# Extreme genetic signatures of local adaptation during plant colonization

**DOI:** 10.1101/485789

**Authors:** Niraj Shah, Tomomi Wakabayashi, Yasuko Kawamura, Cathrine Kiel Skovbjerg, Ming-Zhuo Wang, Yusdar Mustamin, Yoshiko Isomura, Vikas Gupta, Haojie Jin, Terry Mun, Niels Sandal, Fuyuki Azuma, Eigo Fukai, Ümit Seren, Shohei Kusakabe, Yuki Kikuchi, Shogo Nitanda, Takashi Kumaki, Mads Sønderkær, Kaare Lehmann Nielsen, Korbinian Schneeberger, Jens Stougaard, Shusei Sato, Mikkel Heide Schierup, Stig Uggerhøj Andersen

**Author notes:** These authors contributed equally to the work (Niraj Shah, Tomomi Wakabayashi, Yasuko Kawamura). Lead contact Stig U. Andersen, Department of Molecular Biology and Genetics, Aarhus University, Denmark. Authors for correspondence Stig U. Andersen, Department of Molecular Biology and Genetics, Aarhus University, Denmark. Mikkel Heide Schierup, Bioinformatics Research Centre, Aarhus University. Shusei Sato, Graduate School of Life Sciences, Tohoku University, Japan.

## Abstract

Colonization of new habitats is expected to require genetic adaptations to overcome environmental challenges. Here we use full genome re-sequencing and extensive common garden experiments to investigate demographic and selective processes associated with the recent colonization of Japan by *Lotus japonicus*. We carefully track the colonization process where *L. japonicus* gradually spread from subtropical conditions to much colder climates in northern Japan. We characterize the loss of diversity during this process and identify genomic regions with extreme genetic differentiation. Next, we perform population structure-corrected association mapping of phenotypic traits measured in a common garden and discover a number of genome-wide significant associations. Contrasting these analyses, we find that there is a strong concordance between phenotypic variation and extreme differentiation for overwintering and flowering time traits. Our results provide evidence that these traits were direct targets of selection by local adaptation during the colonization process and point to associated candidate genes.

## Introduction

*Lotus japonicus* (*Lotus*) is a perennial wild legume. It has a relatively small genome size of ~472 Mb (Handberg and Stougaard, 1992) and is found in diverse natural habitats across East and Central Asia, including Japan, Korea, and China, and extending west into Afghanistan (Hashiguchi et al., 2011). *L. japonicus* natural diversity is being characterized through the establishment of a collection of wild Japanese accessions (Hashiguchi et al., 2011). They represent an interesting population sample for studying local adaptation because of the geographical isolation of the Japanese archipelago and the pronounced variation in climate between the southern and the northern parts of the country. Climates range from subtropical to hemiboreal with yearly average temperatures of 17.9 and 5.3 °C, respectively, while annual daylight varies from 1316 to 2202 hours, and annual precipitation ranges from 775 mm to 3250 mm (Hashiguchi et al., 2011). The topography of Japan is also very varied, and an up to 3,000 meters high mountain range traverses the central regions of the archipelago. The mountainous topography will likely limit dispersal and the steep climatic gradients could result in strong selection pressures, creating conditions that could promote population differentiation driven by local adaptation (Kawecki and Ebert, 2004).

Experimental evidence for local adaptation can be obtained using reciprocal transplants, which compare at home versus away fitness, or common garden experiments that compare fitness of locals and immigrants in the same environment. Superior fitness at home in reciprocal transplants and locals outperforming immigrants in common gardens constitute evidence of local adaptation, with the local vs. immigrant comparison arguably offering stronger evidence (Kawecki and Ebert, 2004). Such experiments are now being coupled with genotype data to begin understanding the molecular events underlying local adaptation. Once it has been demonstrated that a subpopulation displays local adaptation, genotyping of individuals from contrasting subpopulations allows genome scans for selection signatures such as high fixation index (*F*_ST_) levels. Demographic history, however, can generate similar signatures through genetic drift, causing false positives.

Population structure, whether due to genetic drift or selection, has long been recognized as a major confounding effect in genome-wide association (GWA) studies, and stringent population structure correction methods have been developed to separate genuine genotype-phenotype associations from spurious associations due to population structure (Atwell et al., 2010; Kang et al., 2008). It has been suggested that combining population structure-corrected GWA analysis of phenotypic data from common garden experiments with purely genotype-based genome scans could be a powerful approach for studying local adaptation. The argument is that overlapping adaptive signals detected both using phenotype-independent genome scans and common garden phenotype data constitute independent lines of evidence for local adaptation (de Villemereuil et al., 2016).

Perhaps the most striking example of GWA application to the study of local adaptation is the investigation of human skin pigmentation. Evolution of skin pigmentation is driven by UV irradiation, with dark skin offering protection where there are high levels of UV irradiation, and light skin promoting vitamin D synthesis under low UV irradiation. SNPs in several genes involved in melanin production were identified as associated with skin pigmentation in GWA scans, and they also generally show extremely high *F*_ST_ levels (Crawford et al., 2017). Relatively few studies have applied GWA to the investigation of plant local adaptation. One large *Arabidopsis* study used common garden fitness data in combination with GWA and correlations with climatic factors, but did not employ genome scans for population differentiation (Fournier-Level et al., 2011). A second study examined drought stress data from a laboratory experiment and observed marginally elevated *F*_ST_ levels for the top SNPs (Exposito-Alonso et al., 2018), while a third showed evidence for an adaptive role of a sodium transporter by GWA analysis of ion accumulation coupled with investigation of gene expression and distances to saline soils (Baxter et al., 2010). In addition, flowering time is considered a critical adaptive trait in annual *Arabidopsis* and has been thoroughly examined in GWA studies (1001 Genomes Consortium, 2016; Atwell et al., 2010). However, human dispersal of *Arabidopsis* seeds has eroded the link between geographic and genetic distance, complicating interpretation with respect to local adaptation (1001 Genomes Consortium, 2016; Bergelson et al., 1998; Johanson et al., 2000). Most of the relatively few plant traits studied have thus not revealed clear signals of local adaptation, and the power of combining GWA analysis of common garden phenotype data with genotype-based genome scans for the study of local adaptation has not been systematically explored.

Here, we genetically characterize a set of wild Japanese *Lotus* accessions, identify subpopulations, characterize their demographic history, and show evidence for local adaptation in a common garden experiment. We discover very pronounced overlaps in adaptive signals between phenotype-based GWA and genotype-based *F*_ST_ approaches, allowing us to identify traits and associated genomic loci that were direct targets of selection by local adaptation during *Lotus japonicus* colonization of Japan.

## Results

### Kyushu Island is the center of *Lotus* diversity in Japan

We carried out whole-genome re-sequencing of 136 wild *Lotus* accessions collected throughout Japan using Illumina paired-end reads (**Figure 1A, Supplemental file 1**), identifying a set of 525,800 high confidence SNPs in non-repetitive regions (**Supplemental file 2, Supplemental figures 1-2**).

**Figure 1:**
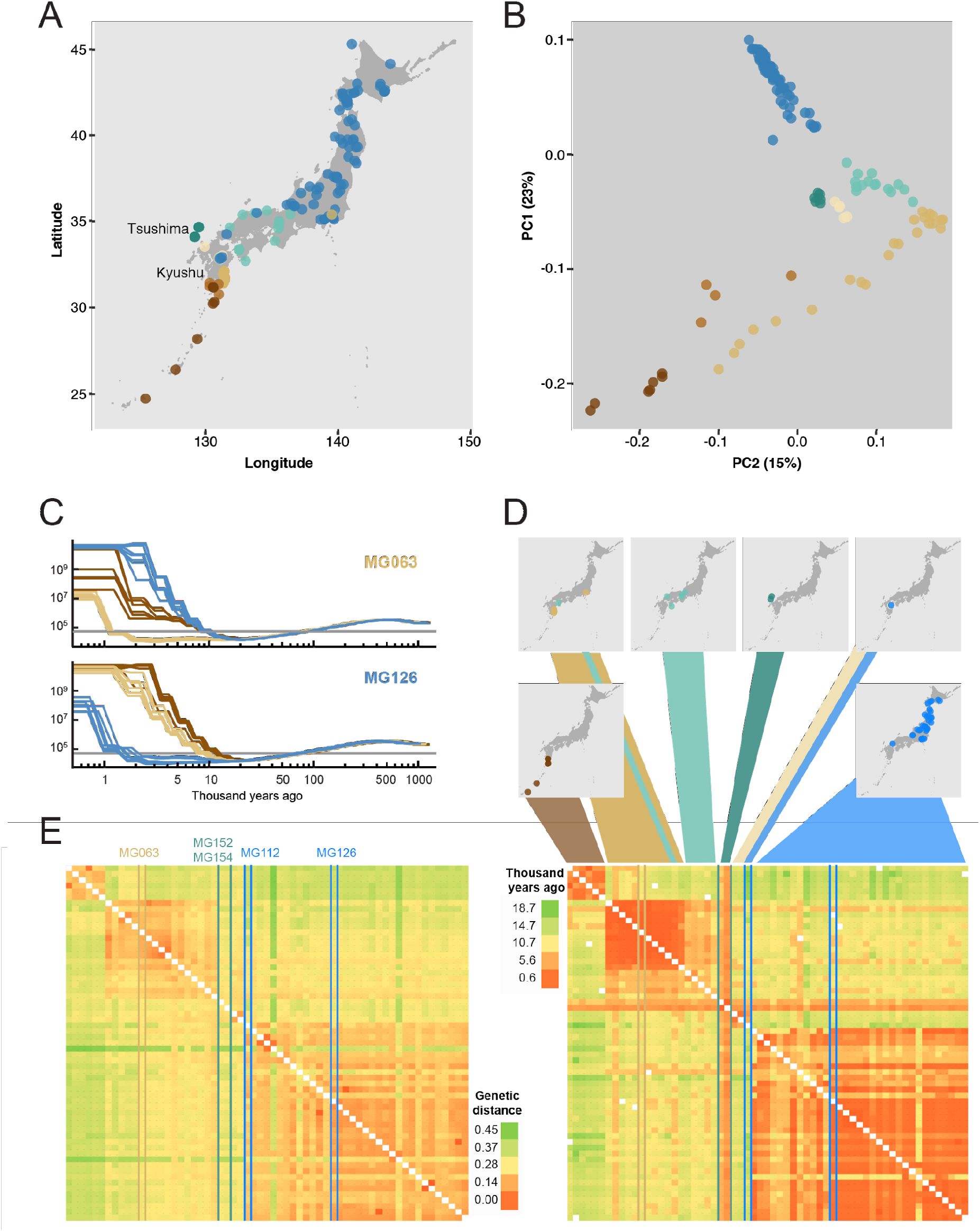
Geography and genetics of Japanese *Lotus* accessions. A) Geographical origin of the *Lotus* accessions. B) Principal component analysis (PCA) based on the genotypes of the *Lotus* accessions. C) Pairwise sequentially Markovian coalescent (PSMC) curves based on pseudodiploids generated by merging alignments of pairs of accessions. Each curve indicates the inferred population size history through time for pseudodiploid pairs including MG063 and MG126, respectively. The horizontal grey line indicates an effective population size of 5×10^4^. D) The lower panel shows a heatmap of divergence times (years ago) for pseudodiploid pairs as estimated from the PSMC analysis with a population size cutoff of 5×10^4^. Accession colors are matched in panels A-E. E) Heatmap showing the genetic distances between pairs of accessions. D-E) Accessions MG152 and MG154 from Tsushima Island (green), MG112 from central Kyushu (blue), MG126 from northern Japan, and M063 from eastern Kyushu are highlighted as indicated in E). The accessions are shown in the same order in panels D) and E).

In order to examine the genetic relatedness of the accessions, we carried out a principal component (PCA) analysis based on the accession genotypes. We found a striking correlation between the position of the accessions in the PCA plot and their geographical origin (**Figure 1A-B**), similar to that observed for studies of human populations (Bryc et al., 2010; Novembre et al., 2008). PC1 clearly separated the north and south of Japan, while PC2 most strongly distinguished accessions from the eastern and southern coastline of Kyushu Island (**Figure 1A-B**). For both PC1 and PC2, the northern accessions clustered more tightly than the southern accessions, and the accessions from Kyushu Island were particularly well-resolved (**Figure 1A-B**).

The tight clustering of the northern accessions suggested that these could have lower levels of genetic diversity and, consequently, that *Lotus* could have arrived in the south of Japan and then migrated north. To examine the migration history of *Lotus* in detail, we carried out a Pairwise Sequentially Markovian Coalescent (PSMC) model analysis, which infers population size history from a diploid sequence based on the extent of heterozygous regions in the genome (Li and Durbin, 2011). Since *Lotus* is self-compatible and most accessions had low heterozygosity levels (**Supplemental file 1**), the individual accessions were not well suited for PSMC analysis. Instead, we generated pseudo-diploids by merging read alignments from pairs of a subset of individuals with at least 7x read coverage and called a pseudo-diploid consensus sequence based on the merged alignments, which was used for PSMC analysis. This type of analysis can infer the time of last contact of the pair of individuals forming the pseudo-diploid as the time of the last coalescence events, measured as an abrupt increase in the estimated effective population size. We exploited this feature to roughly estimate divergence times (cessation of gene flow) for the pseudodiploid pairs as the point in time when the PSMC curve abruptly rises (**Figure 1C-D**), assuming a mutation rate of 6.5×10-9 per year (Ossowski et al., 2010). For example, for accession MG126 found in northern Japan, the cessation of gene flow to other accessions from northern Japan (blue lines) are 1-5 thousand years, for Eastern Kyushu (light brown lines) 7-9 thousand years and for south and west Kyushu (dark brown lines) 10-12 thousand years (**Figure 1C-D**). The estimated divergence times for all pairs of accessions showed clear clustering along the inferred colonization route (**Figure 1D**) with South and Northern Japan having last contact 10-18 thousand years ago, consistent with colonization after the last ice age. For the same accession pairs, we also calculated simple genetic distances (**Figure 1E**). Despite the clear overall similarity of the genetic distance and PSMC-based divergence time analysis, we found marked deviations for accessions from Tsushima Island (MG152 and MG154) and central Kyushu (MG112). The Tsushima accessions appeared to have diverged recently from all other accessions (**Figure 1D**), but the genetic distance analysis indicated relatively large genetic distances from the Tsushima lines to all other accessions with no strong association to any other population cluster (**Figure 1E**). This unique signature and the central location of these accessions in the PCA plot (**Figure 1B**), suggests that Tsushima Island may represent a point of origin for all of the Japanese *Lotus* populations. Among the central Kyushu lines, MG112 is the southernmost accession belonging to blue cluster in the PCA plot (**Figure 1A,D**). In the genetic distance analysis, MG112 is clearly most similar to the other accessions from this PCA cluster, but it shows long divergence times from these and shorter divergence times from its fellow central Kyushu lines (**Figure 1D-E**). Likewise, MG110 and MG111 show long divergence times from other accessions in their geographical vicinity (except for the Tsushima lines), but shorter divergence times with the central Kyushu lines (**Figure 1D-E**). These central Kyushu lines, which grow at high altitudes with sub-0°C minimum temperatures, thus likely became isolated and stopped exchanging genetic material with other accessions relatively shortly after *Lotus* colonisation of Japan. Altogether, our data suggests that *Lotus* migrated north and south from a starting point near Tsushima Island and that the north was colonized most recently, since accession pairs across a large geographical area in northern Japan display relatively short divergence times (**Figure 1D and Supplemental file 1**). Our results also indicate that representatives of all major genetic clusters are found on Kyushu Island, making it the center of diversity for *Lotus* in Japan.

### The Japanese *Lotus* accessions can be grouped into three subpopulations

In the PCA analysis, the northern and southern accessions clustered into different groups, indicating that distinct subpopulations might exist. fastSTRUCTURE (Raj et al., 2014) analysis, which groups accessions based on allele frequencies, suggested that a large fraction of the variation could be accounted for by three subpopulations (**Supplemental figure 3**). Populations 1 (pop1), 2 (pop2), and 3 (pop3) occupied southern, central, and northern Japan, respectively, and corresponded to the three vertices in the PCA plot (**Figure 2A-C**). The three subpopulations also overlapped with the major clusters identified in the analysis of pairwise genetic distances and divergence times (**Figure 1D-E**), indicating that they reflect a robust grouping of the accessions. This is supported by Figure 2D, which shows that the genetic distance for a given physical distance is larger for comparisons between these three populations as compared to distances within populations. Examining genetic diversity (pi) for each subpopulation, we found that the southernmost populations showed higher levels of diversity (**Figure 2E**), which is consistent with the hypothesis that *Lotus* migrated north from southern Japan, losing genetic diversity along the way.

**Figure 2.**
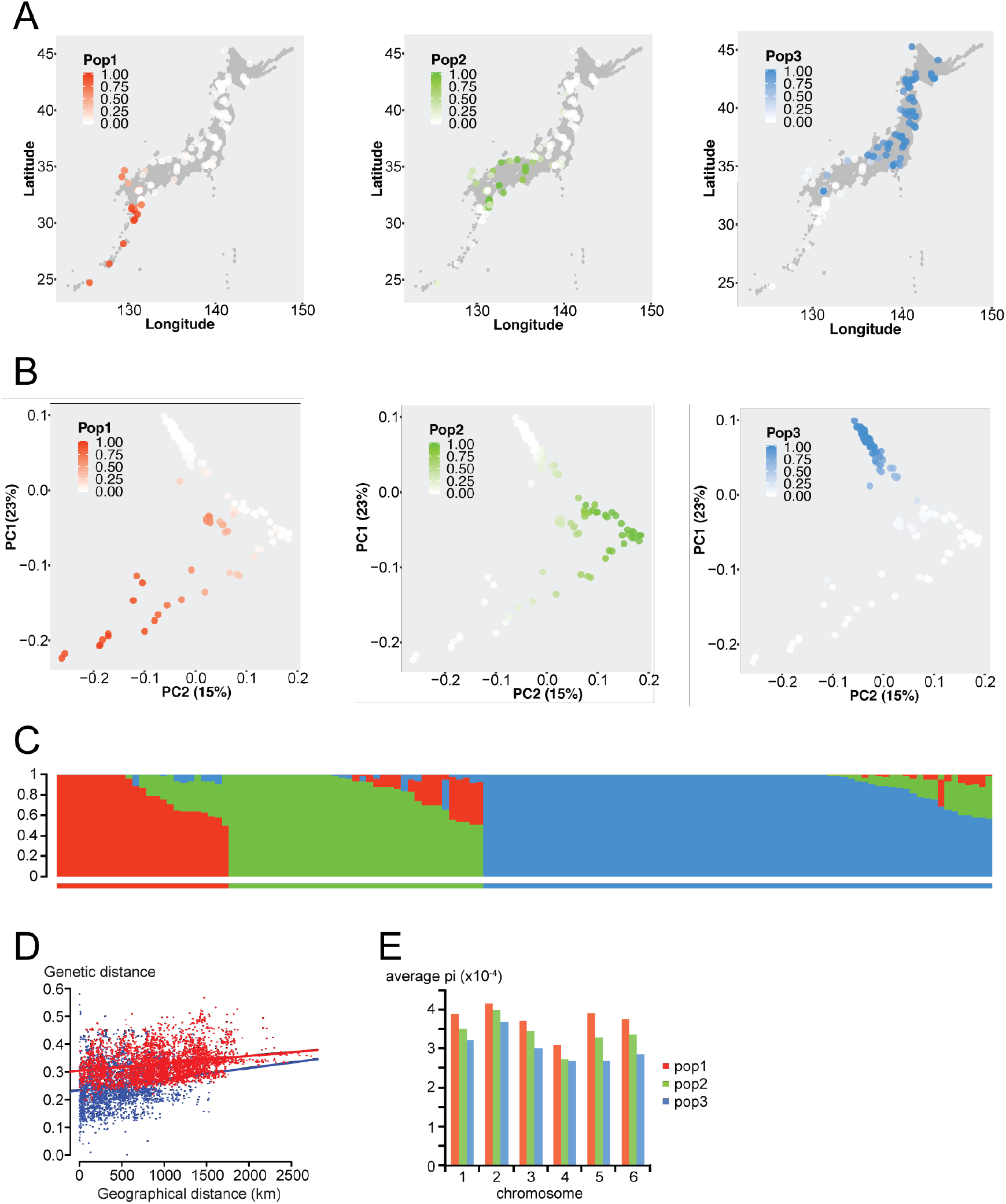
Pairwise divergence times and genetic distances. **A-B)** Accessions are colored by their population membership placed according to geographical origin (A) or according to their coordinates in the PCA analysis (B). **C)** STRUCTURE plot showing population memberships for pop1 (red), pop2 (green) and pop3 (blue). **D)** Genetic distance versus geographical distance. Red: between-population comparisons. Blue: within population comparisons. **E)** Average nucleotide diversity (pi) per chromosome.)

### Contrasting subpopulations identifies strong population differentiation signals

We calculated *F*_ST_ for all polymorphic positions, comparing pop3 accessions against accessions with no population 3 membership in order to detect markers strongly differentiated between these two groups with relatively many characterized members. There were 5,612 genes with at least four informative SNPs, for which we calculated the mean *F*_ST_ per gene. The median per-gene *F*_ST_ value was 0.24, and there were 132 genes with average *F*_ST_>0.65 (**Supplemental figure 4A-B**). The top gene *Lj6g3v1790920*, had a mean *F*_ST_ value of 0.99 across 20 SNPs and was located in a ~20 kb genomic region showing very strong fixation of the alternative allele in pop3 relative to accessions without pop3 membership (**Supplemental figure 4C**).

### Pop3 is locally adapted

The distinct regions occupied by each of the three subpopulations, the restriction of pop3 accessions in southern Japan to a cold, high-altitude environment in central Kyushu, and the pronounced differentiation observed for specific genomic regions, suggested that population differentiation could have been influenced by local adaptation in addition to genetic drift. To test this hypothesis experimentally, we grew the accessions in a common garden using a field site at Tohoku University (38.46°N, 141.09°E), which is located in northern Japan where pop3 individuals dominate (**Figure 2A and Figure 3A**). A number of plants died during winter, providing us with a direct fitness measure, and the accession overwintering rate showed a strong correlation with geographic origin (**Figure 3B-C, Supplemental figures 5-6**). To test for evidence of local adaptation of pop3 to its native northern Japan environment, we compared winter survival as a function of population membership for non-admixed accessions. Across all three years, pop3 accessions clearly outperformed both pop1 and pop2 individuals in terms of survival (P<1e-4, generalized linear model ANOVA with Tukey multiple comparison test), providing evidence of pop3 local adaptation (**Figure 3B**).

**Figure 3.**
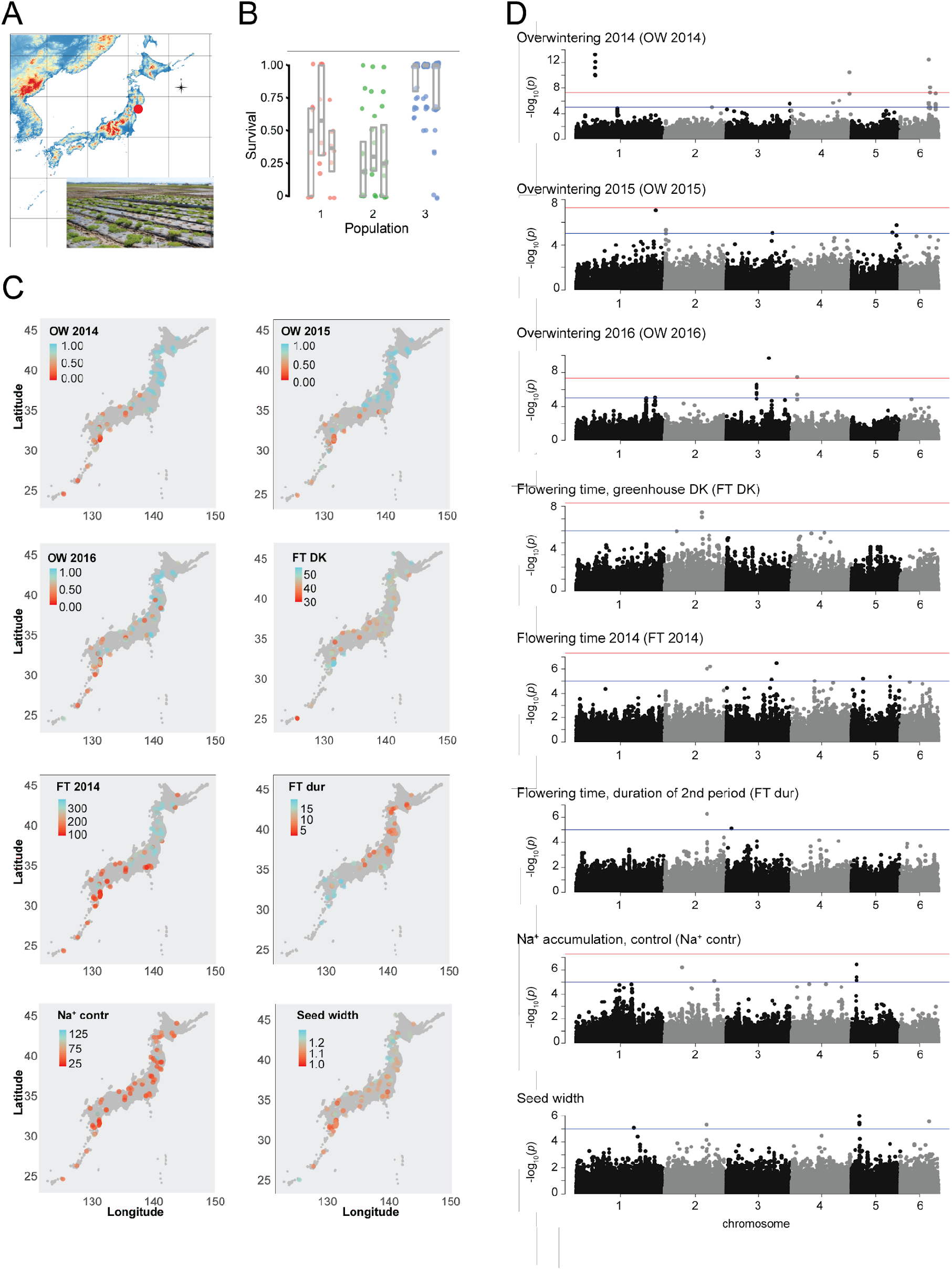
Phenotype data and GWA analysis. **A)** The site in Kashimadai, 38.46 N, 141.09 E, where the field experiments were carried out is denoted by a red circle. **B)** Boxplot showing winter survival for non-admixed individuals from pop1-3. For each population, the three plots show data from 2014, 2015 and 2016, respectively. The horizontal grey bar indicates the median, and the lower and upper hinges correspond to the 25th and 75th percentiles. **C)** Heatmaps displaying phenotype data. OW: overwintering, the fraction of plants surviving winter. FT: flowering time. FT DK: days to flowering in a greenhouse located at 56.229406N, 10.126424E, 8380, Trige, Denmark. FT 2014: days to first flowering at the Kashimadai field site in 2014. FT dur: duration of the second flowering time period at the Kashimadai field site in 2014/2015. Na^+^ contr: sodium ion accumulation in roots. **D)** Manhattan plots showing results of GWA scans for the phenotypes indicated. Blue line: *p* = 10^-5^. Red line: *p*=10^-7^.

We also collected phenotype data for other potentially adaptive traits. Flowering time was a likely candidate, and to maximize the chances of identifying phenotypic variation of adaptive relevance, we carried out both a field and a greenhouse flowering time experiment. In the field, we found the flowering time characteristics to depend greatly on the planting date. In 2014, the plants were sown on July 4th and transplanted to the field on August 4th. This late planting date caused a very strong differentiation between the accessions. Many of the northern accessions failed to flower in the planting year, flowering instead the following spring, and this trait was strongly correlated with geographic origin, in contrast to the greenhouse flowering time data (**Figure 3C, Supplemental figures 5-6**). In the field, we also measured leaf accumulation of sodium and potassium ions, which was poorly correlated with geographic origin (**Figure 3C, Supplemental figures 5-6**). In addition, we quantified seed properties, which showed an intermediate correlation with geographic origin (**Figure 3C, Supplemental figures 5-6**).

### Selection acted on winter survival during pop3 local adaptation

For our population sample, linkage disequilibrium decayed to an *r^2^* value of 0.2 within approximately 10 kb, indicating that mapping resolution would be sufficiently high to identify a limited sets of candidate genes in GWA scans (**Supplemental figure 7**). With phenotype data available for potentially adaptive traits, we proceeded with GWA analysis using a method that includes stringent correction for population structure, ensuring that significant hits represent associations that cannot be explained by the overall population structure (Atwell et al., 2010; Kang et al., 2008). We identified SNPs strongly associated with a number of the examined traits, particularly for overwintering and flowering time (**Figure 3D**). Overwintering 2014 displayed the most highly significant associations, which were distributed across several separate genomic loci, indicating that winter survival is a complex trait. The 57 top SNPs (P<5×10^-5^) explained 48% of the phenotypic variation for overwintering 2014.

We then compared the GWA and *F*_ST_ results to check for skews in the *F*_ST_ value distribution for the top GWA SNPs. The SNPs associated with most flowering time traits as well as the ion accumulation traits did not show large deviations from the overall distribution of *F*_ST_ values (**Figure 4A-B, Supplemental figure 8**). However, SNPs associated with overwintering traits, flowering time in 2014, the duration of the second flowering time period, and seed width showed an enrichment for high *F*_ST_ values (**Figure 4A-B, Supplemental figure 8**). Moreover, it was characteristic for these traits that even among the SNPs associated with the phenotype (using a *P*-value threshold of 10^-3^), it was the most strongly associated SNPs that showed the highest *F*_ST_ values (**Figure 4A-B rightmost panels, Supplemental figure 8**). This was in contrast to for instance greenhouse flowering time, which also showed a significant *F*_ST_ skew for the GWA SNPs, but where the top GWA hits showed only intermediate *F*_ST_ values (**Figure 4A-B, Supplemental figure 8**).

**Figure 4.**
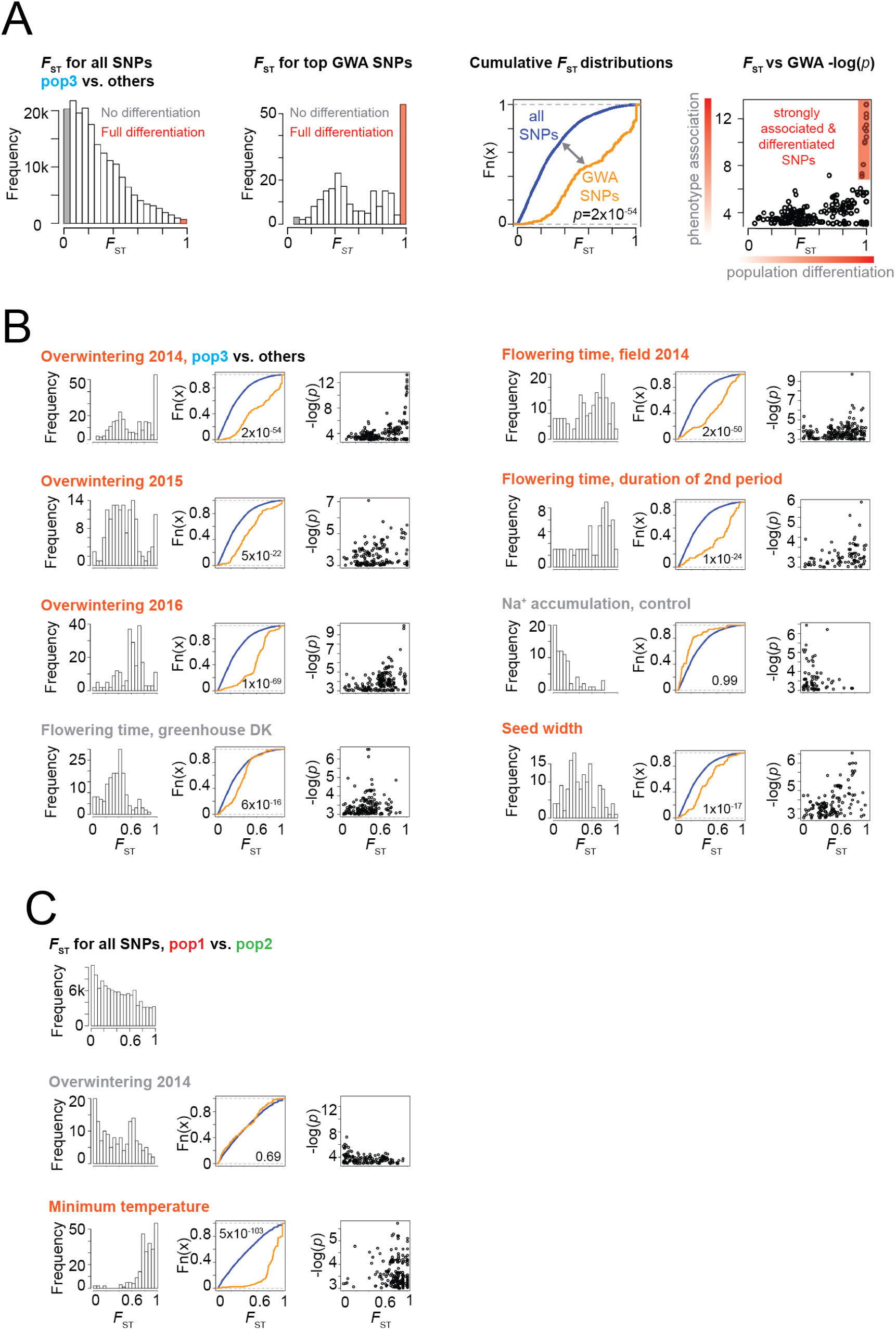
Distribution of *F*_ST_ values for top GWA hits. **A)** Reading guide for panels B) and C). **B-C)** The leftmost panel shows distributions of *F*_ST_ values for the SNPs with -log(*p*) GWA scores > 3 for the trait indicated. The middle panel shows experimental cumulative distributions graphs for the *F*_ST_ values for all SNPs (blue) and the SNPs shown in the leftmost panel (orange). If the orange curve lies below the blue curve, it indicates a shift to higher *F*_ST_ values for the SNPs with high GWA scores. The *p*-value that the blue curve lies below the orange curve is indicated (Kolmogorov-Smirnov test). The rightmost panel shows *F*_ST_ values plotted versus GWA -log(*p*) scores. **A,B)** *F*_ST_ values were calculated based on a pop3 vs. non-pop3 comparison. **C)** *F*_ST_ values were calculated based on a pop1 vs. pop2 comparison. Traits likely associated with local adaptation are highlighted (orange).

Whereas pop3 was locally adapted to the field site near Tohoku, the Tohoku winter survival phenotype should not be associated with pop1 vs. pop2 differentiation. To test this, we repeated the GWA/*F*_ST_ overlap analysis for the pop1 vs. pop2 comparison. As expected, the SNPs associated with Tohoku overwintering traits did not show skews towards high *F*_ST_ values for the pop1 vs. pop2 comparison (**Figure 4C, Supplemental figure 9**). In contrast, the top SNPs associated with minimum temperature at the geographic origin of the accessions showed a very strong skew towards high *F*_ST_ values for pop1 vs pop2 differentiation (**Figure 4C, Supplemental figure 9, Supplemental file 3**), suggesting that this environmental variable was likely critical for pop1/pop2 differentiation.

As a visual illustration, we overlaid GWA and *F*_ST_ signals for the traits and chromosomes showing the strongest GWA hits (**Figure 5, Supplemental files 4-5**). The most striking example of overlap between GWA and *F*_ST_ signals was found for overwintering 2014 on chromosome 6, where the top GWA hit was found in the region containing the genome-wide top *F*_ST_ gene *Lj6g3v1790920* (**Supplemental figure 4C**), and two additional strong GWA signals overlapped with prominent *F*_ST_ peaks (**Figure 5**). The top 2014 overwintering GWA signal was also recovered in 2015, albeit with lower signal strength (**Figure 5**). The overlap between the top GWA hits for overwintering and the genome-wide top *F*_ST_ signals indicates that winter survival has played a major role in pop3 local adaptation and points to specific candidate genes with potential adaptive roles.

**Figure 5.**
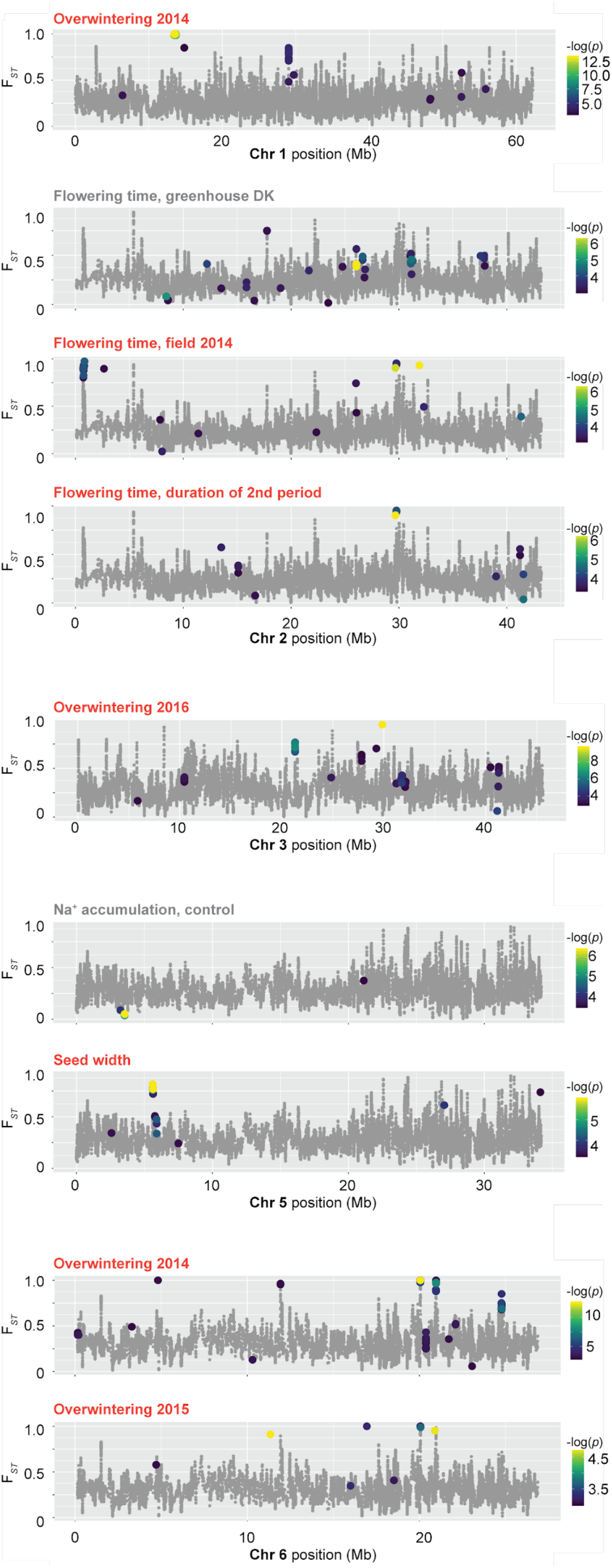
Overlay of *F*_ST_ values and GWA results. *F*_ST_ averages for 10 SNPs (grey dots) are plotted. Colored dots show *F*_ST_ values for SNPs with GWA -log(*p*) scores > 3 for the chromosomes and traits indicated.

Three candidate genes reside within the strongly differentiated chr6 region 20,048,000-20,068,000 comprising the top GWA overwintering signals. *Lj6g3v1790920* encodes a RING-type zinc finger protein, *Lj6g3v1790910* is a ubiquitin ligase, while *Lj6g3v1789900* is a putative 1-aminocyclopropane-1-carboxylate oxidase involved in ethylene biosynthesis. Another prominent GWA peak on chr6 was in the region 20,961,109-20,970,404, containing a single candidate gene, *Lj6g3v1887780*, which is annotated as an HBS1-like protein containing a predicted Alpha/Beta hydrolase fold (IPR029058). The third prominent GWA peak on chr6, 24,760,760-24,781,399 contained three candidate genes, two putative G-type lectin S-receptor-like serine/threonine-protein kinases, *Lj6g3v2130140* and *Lj6g3v2130160*, and *Lj6g3v2130130* encoding a putative Glutamyl-tRNA(Gln) amidotransferase subunit A protein. On chr1, several SNPs with very strong GWA associations and high *F*_ST_ values were dispersed across a large 240 kb region (13,458,027-13,698,337). A more clearly defined peak on chr1 was located at 29,029,933-29,033,958. It contained a single candidate gene, *Lj1g3v2533770*, which encodes a FERONIA receptor-like kinase. According to published expression data, the overwintering candidate genes were primarily expressed in roots and nodules (**Supplemental figure 10**) (Høgslund et al., 2009; Mun et al., 2016; Verdier et al., 2013).

### Flowering time and seed properties may have contributed to pop3 local adaptation

Although greenhouse flowering time produced signals in the GWA analysis (**Figure 3D**), the SNPs identified did not show a pronounced skew towards high *F*_ST_ values (**Figure 4B**), and the trait was poorly correlated with geographical origin (**Supplemental figures 5-6**). In contrast, the SNPs strongly associated with flowering time field data from 2014, where the accessions were planted late and showed strong differentiation, did overlap with high *F*_ST_ scores, as did the duration of the 2nd flowering time period (**Figure 4B**). Remarkably, the top GWA hits on chromosome 2 overlapped between 2014 flowering time and duration of the second flowering time period (**Figure 5**), suggesting that the same gene(s) could be involved in controlling both the onset of flowering and the duration of the flowering time period. The genes, *Lj2g3v1988990* and *Lj2g3v1989150*, neighboring the top GWA hits are expressed in flowers, pods, and seeds (**Supplemental figure 10**) (Høgslund et al., 2009; Mun et al., 2016; Verdier et al., 2013). *Lj2g3v1988990* is a putative glutamate dehydrogenase, while *Lj2g3v1989150* is highly similar to *AtEMF2* (*AT5G51230*) and *AtVRN2* (*AT4G16845*), which are both flowering time regulators, with *VRN2* mediating vernalization responses in Arabidopsis (Chen et al., 1997; Gendall et al., 2001).

Seed width, which showed less strong correlations with environmental parameters and geographical origin (**Figure 3C, Supplemental figures 5-6**), also showed GWA hits with a skew towards high *F*_ST_ values, although not as extreme as for the overwintering or flowering time traits (**Figure 4B**). This was due to a peak on chromosome 5 (**Figure 5**), where the genes *Lj5g3v0526490* and *Lj5g3v0526510*, highly expressed in seeds and flowers (**Supplemental figure 10**) (Høgslund et al., 2009; Mun et al., 2016; Verdier et al., 2013), but without known functions, neighboured the top scoring SNPs.

## Discussion

Here, we have inferred the population structure and demographic history of wild *Lotus* in Japan. We conclude that Lotus colonized Japan from Tsushima Island after the last ice ages in a very regular fashion, reaching southern Japan before the northern part with little long-distance gene flow post colonization. It was *a priori* conceivable that the very different climate across Japan could be a barrier to colonization and that selection on standing variation for phenotypic traits related to temperature and daylight would be detectable. Thus, we have exploited the *Lotus* subpopulations to study local adaptation through a combination of genotype-based *F*_ST_ genome scans and phenotype-based GWA analyses using data from a common garden field experiment. Our study illustrates the power of this combination. Alone, *F*_ST_ genome scans identified strong population differentiation signals, but provide no hints as to the phenotypic traits or environmental variables driving the differentiation, nor do they account for the effects of population structure. GWA analysis of phenotypic traits, on the other hand, provided strong, population structure-corrected phenotype-genotype associations for a number of traits, but offered no way of determining which, if any, of the traits were important for local adaptation. In our study, the combination of the two approaches was powerful, because we found very pronounced skews towards high *F*_ST_ values for certain traits, identifying these as strong candidates for driving local adaption and supplying trait labels to previously anonymous *F*_ST_ peaks.

The year-to-year differences in winter survival and their impact on the GWA results illustrates that the conditions under which the common garden experiments are carried out have a large impact on the power to detect overlapping signals from GWA and *F*_ST_ genome scans. In our case, the top overwintering GWA signals overlapped most clearly with the top *F*_ST_ peaks for winter 2014. Here, the plants were transplanted late, which resulted in very strong differentiation between adapted and non-adapted accessions. Subsequent years, planting took place earlier in the year to promote plant survival and allow quantification of other phenotypic traits, resulting in less pronounced GWA signals, and when the full experiment was re-sown in 2017, the earlier sowing date and milder winter resulted in survival of nearly all accessions, thwarting GWA analysis.

These observations also offer a possible explanation for why similar *F*_ST_/GWA overlaps have not been frequently reported, since their identification might require screening several candidate phenotypic traits of putative adaptive relevance for the subpopulations under investigation, rather than for instance focusing on a single trait phenotyped across many individuals and subpopulations (Exposito-Alonso et al., 2018). On the same note, we found it interesting that the top GWA SNPs associated with greenhouse flowering did not show a skew towards high *F*_ST_ values, but that the SNPs associated with flowering time in 2014 and flowering time duration of second year plants did show pronounced skews (**Figure 4B**) and pointed to a putative ortholog of an *Arabidopsis* flowering time regulator. The flowering time data again emphasized the impact of genotype by environment interactions and the need to screen multiple phenotypes to identify traits and associated genes of relevance for local adaptation.

For a perennial plant such as *Lotus*, it makes intuitive sense that surviving harsh winters could have been critical for local adaptation to cold climates. Our work supported this hypothesis and identified the most prominent genomic regions and candidate genes associated with overwintering. Remarkably, 48% of the phenotypic variation for winter survival 2014 could be explained by strongly differentiated and phenotype-associated SNPs, indicating that a limited number of loci with large effects have a strong impact on the trait. Such pronounced overlaps between top GWA and *F*_ST_ signals have not, to our knowledge, been reported in the plant literature, but they are reminiscent of the characteristics displayed by SNPs associated with human skin pigmentation (Crawford et al., 2017). Analogously to the tradeoff between UV protection and vitamin D synthesis, our results suggest that strong divergent selection is acting on *Lotus* alleles that are beneficial for perennial winter survival in cold climates, but likely have detrimental effects on plant fitness in warmer regions, thus contributing to shaping and maintaining genetically differentiated populations that are locally adapted to markedly different climates. Based on the candidate genes we identify here, the molecular genetic mechanisms that underpin these fundamental adaptations can now be investigated.

## Materials and methods

### Sequencing, mapping and variant calling

All samples were sequenced using Illumina paired-end reads (ENA accession PRJEB27969). The reads were mapped to the *L. japonicus* genome version 3.0 (http://www.kazusa.or.jp/lotus/) using Burrows-Wheeler Aligner (BWA) *mem* v0.75a with default parameters (Li, 2013) (**Supplemental file 6**). An average of 96% of the reads were mapped and the resulting average genome coverage was 12.5 (**Supplemental file 1**). Duplicates were marked in the resulting alignment (BAM) files using Picard v1.96. Then, using the Genome Analysis Tool Kit (GATK) v2.7-2 pipeline (McKenna et al., 2010), reads were re-aligned in INDEL regions using the *RealignerTargetCreator* and *IndelRealigner* functions. A preliminary list of SNPs and INDELs was then generated using the re-aligned BAM files using the GATK *UnifiedGenotyper* function. Subsequently, base recalibration of the scores was done using the *BaseRecalibrator*, and the BAM files generated were compressed using the *ReduceReads* function followed by variant discovery, resulting in 8,716,552 putative variants that included SNPs (7,836,221) and INDELs (880,331). See **Supplemental file 6** for details.

The highly inbred reference accession, MG-20, was also re-sequenced. The polymorphic positions called as homozygous reference and heterozygous in MG-20 were compared to the positions with MG-20 homozygous reference calls, and the properties of these positions were analyzed. Low inbreeding coefficient and high haplotype scores characterized heterozygous positions in MG-20 (**Supplemental figure 1**). The INDEL positions had similar properties. The SNP positions were therefore required to fulfill the following criteria: Inbreeding Coefficient > 0.1, HaplotypeScore < 0.3, alternative allele quality > 30, total depth > 150 and a homozygous reference call for the MG-20 accession. Callable positions were defined as positions that fulfilled the above-mentioned criteria and had a genotype call for at least one accession in addition to MG-20.

To validate our filtering, we compared these variants to a set of independently validated SNPs genotyped using the Illumina GoldenGate system. Out of 343 validated SNPs included in the initial call set, 324 matched the genotype calls based on Gifu B-129 and MG-20 sequencing reads (**Supplemental file 1**). Our stringent filtering had removed 93% of the initially called SNPs and 37% of the experimentally validated SNPs, resulting in a 9.5 fold enrichment for validated SNPs in the filtered set. Assuming that, since *Lotus* Gifu is highly inbred, all heterozygote calls in this accession would be false positives, we estimated a false positive rate of 4%. The site frequency spectrum for the high-confidence SNPs indicated that we had undercalled rare heterozygous alleles with our conservative filtering approach, most likely because these calls would only be supported by data from a small subset of the accessions (**Supplemental figure 2**).

### Calculation of genetic diversity, genetic and geographical distances

The --window-pi function of VCF tools version 0.1.9 (Danecek et al., 2011) was used to calculate genetic diversity, with the chromosome size as the window size (**Supplemental file 6**). Pairwise genetic distances were calculated by assigning a score of +2 for homozygous differences, +1 for heterozygous differences and 0 for identical alleles at each polymorphic site and then dividing by the total number of polymorphic sites. Geographical distances were calculated using the distVincentyEllipsoid function from the R package geosphere. See https://github.com/ShahNiraj/JapanHistory.

### Principal component and linkage disequilibrium analysis

PCA analysis was performed with EIGENSOFT v6.0beta (Price et al., 2006; Pritchard et al., 2000) using the *smartpca* command with default parameters (**Supplemental file 6**). PCA plots were generated using R version 3.4.3 (**Supplemental file 7**). Linkage disequilibrium (LD) was calculated using VCFtools v0.1.9 (Danecek et al., 2011) for all the SNP-pairs with a maximum physical separation of 50,000 base pairs. Chromosome 0 was ignored in the analysis. A randomly selected set of 200,000 SNP-pairs was plotted using R version 3.4.3.

### Population structure analysis

To infer population structure, a Bayesian model based clustering analysis was conducted with fastSTRUCTURE version 1.0 (Raj et al., 2014) for 201,694 non-repetitive SNP markers with minor allele count > 5 in 136 accessions. The analysis was run with a number of clusters (K) ranging from 1 to 8 with default parameter settings.

### PSMC analysis

For estimation of population size history, the Pairwise Sequentially Markovian Coalescent (PSMC) model was used (Li and Durbin, 2011). Accessions with more than 10x coverage were subsampled to 10x coverage. Diploid consensus sequences were made with SAMtools v1.3 and BCFtools v1.3 and SNPs which had mapping quality >50 and read depth between 5 and 100 were included in the analysis. PSMC analysis was run with each pseudodiploid consensus sequence using the recommended settings (Li and Durbin, 2011). The mutation rate per nucleotide in *A. thaliana*, 6.5 × 10^-9^ year/site (Ossowski et al., 2010) was used in the visualization step. See **Supplemental file 6** for the scripts and commands used.

### Phenotyping

To investigate the plant phenotypes under field conditions, we grew the wild *Lotus* accessions in a field at Kashimadai, Graduate School of Life Sciences, Tohoku University (38.46°N, 141.09°E) located in Miyagi Prefecture, Japan from 2014. Partly scrubbed seeds were sown on cell trays containing a 1:1 mixture of soil and vermiculite, and grown in a greenhouse for around one month before transplanting to the field. Sowing and transplanting dates were 3rd of July and 4th of August in 2014, 30th of May and 30th of June in 2015, 3rd of June and 13th of July in 2016, and 30th of March and 28th of April in 2017, respectively. From 2014 to 2017, a total of 106 wild accessions were grown in the same field, including overwintered individuals in 2015 to 2017 seasons. Three individuals were planted in a single spot with 60 cm intervals and each line assigned to two spots each for control and salt stressed fields. Salt stress treatments were conducted by irrigation with underground water containing salt around 1/4 concentration of seawater and normal water (Control). Leaf samples from three plants of each line were taken after four weeks of treatment for measurement of ion contents (Na^+^ and K^+^) in 2014. In the 2017 season, a new field was used for growing 136 wild accessions in control field only.

Greenhouse flowering time was quantified as days to the first open flower in a greenhouse located at 56.229406N, 10.126424E, 8380, Trige, Denmark. Seeds were germinated on wet filter paper and seedlings were transplanted to the greenhouse on May 10th, 2014, where they were grown in peat (Pindstrup, mix 2) mixed with 20% perlite at 18-23 °C (day) 15°C (night) using a 16/8 hour day/night cycle and 70% relative humidity fertilized with Pioner NPK Makro 14-3-23 +Mg blå.

For seed phenotyping, seeds were collected from the plants grown in a greenhouse (Trige Aarhus, Denmark) in 2014. Seed images were obtained using Epson Perfection 600 Photo scanners. The image files were analyzed using SmartGrain (Tanabata et al., 2012) to count total number of the seeds and measure area size, length, width, length-width ratio, circularity of each seed.

### *F*_ST_ and GWA analysis

The --weir-fst-pop function from VCF tools version 0.1.9 (Danecek et al., 2011) was used to calculate fixation indices (*F*_ST_) (**Supplemental file 6**). Prior to GWA analysis, missing genotype data points were imputed using Beagle version 5.0 (Browning et al., 2018), and the VCF file was converted to a simple genotype-only format (**Supplemental files 6 and 8**). GWA scans were carried out using a python implementation of EMMA (https://app.assembla.com/spaces/atgwas/git/source) (Atwell et al., 2010; Kang et al., 2008) with a minor allele count cutoff of 8 (**Supplemental file 6**). The software GCTA-GREML was used to estimate the proportion of phenotypic variance explained by significant SNPs. For selected traits, the method was applied to capture variance explained by a subset of associated SNPs GWA *p*-values≤5e-5. SNPs with a minor allele count below 10 were not included.

## Data availability

Accessions read data have been deposited in the European Nucleotide Archive (ENA) database under primary accession number PRJEB27969 and secondary accession number ERP110110. Genotype data is available for online GWA analysis through a website (https://lotus.au.dk/gwas/) based on the GWAPP platform (Seren et al., 2012). The phenotype data and GWA analyses are also available from the site (https://lotus.au.dk/gwas/#/study/68/overview). Custom scripts and workflows are available at https://github.com/ShahNiraj/JapanHistory and in **Supplemental files 6-7**.

## Author contributions

Conceptualization, S.U.A., S.S., and M.H.S.; Methodology, S.U.A., M.H.S., and S.S.; Software, Ü.S.; Validation, N.Sh., S.U.A., M.H.S, and S.S.; Formal Analysis, N.Sh., T.W., S.U.A., V.G., C.K.S, E.F., and K.S.; Investigation, Y.Ka., M-Z. W., M. S., Y. I., H.J., T.M., N.Sa., S.K., Y. Ki., S. N., T. K. and M.S.; Resources, J.S., S.S., M.H.S, and S.U.A.; Data Curation, N.Sh.; Writing – Original Draft, S.U.A. and N.Sh.; Writing – Review & Editing, S.U.A. and M.H.S.; Visualization, N.Sh., T.W., S.U.A., and C.K.S.; Supervision, S.U.A., S.S., and M.H.S. Project Administration, J.S., S.S., and S.U.A.; Funding Acquisition, J.S., S.S., S.U.A., and M.H.S.

## Supporting information

## Acknowledgements

The work was supported by the Danish National Research Foundation grant DNRF79 (JS), the Genome Information Upgrading Program of the National BioResource Project in 2014 (SS), a JST CREST grant (number JPMJCR16O1) (SS), grant no 6108-00385A from The Danish Council for Independent Research | Natural Sciences (MHS) and grant no. 10-081677 from The Danish Council for Independent Research | Technology and Production Sciences (SUA). Wild accessions of *L. japonicus* were provided by the National BioResource Project ‘Lotus/Glycine’. We wish to thank Thomas Bataillon for critical reading of the manuscript.

## Supplemental figures

**Supplemental figure 1:**
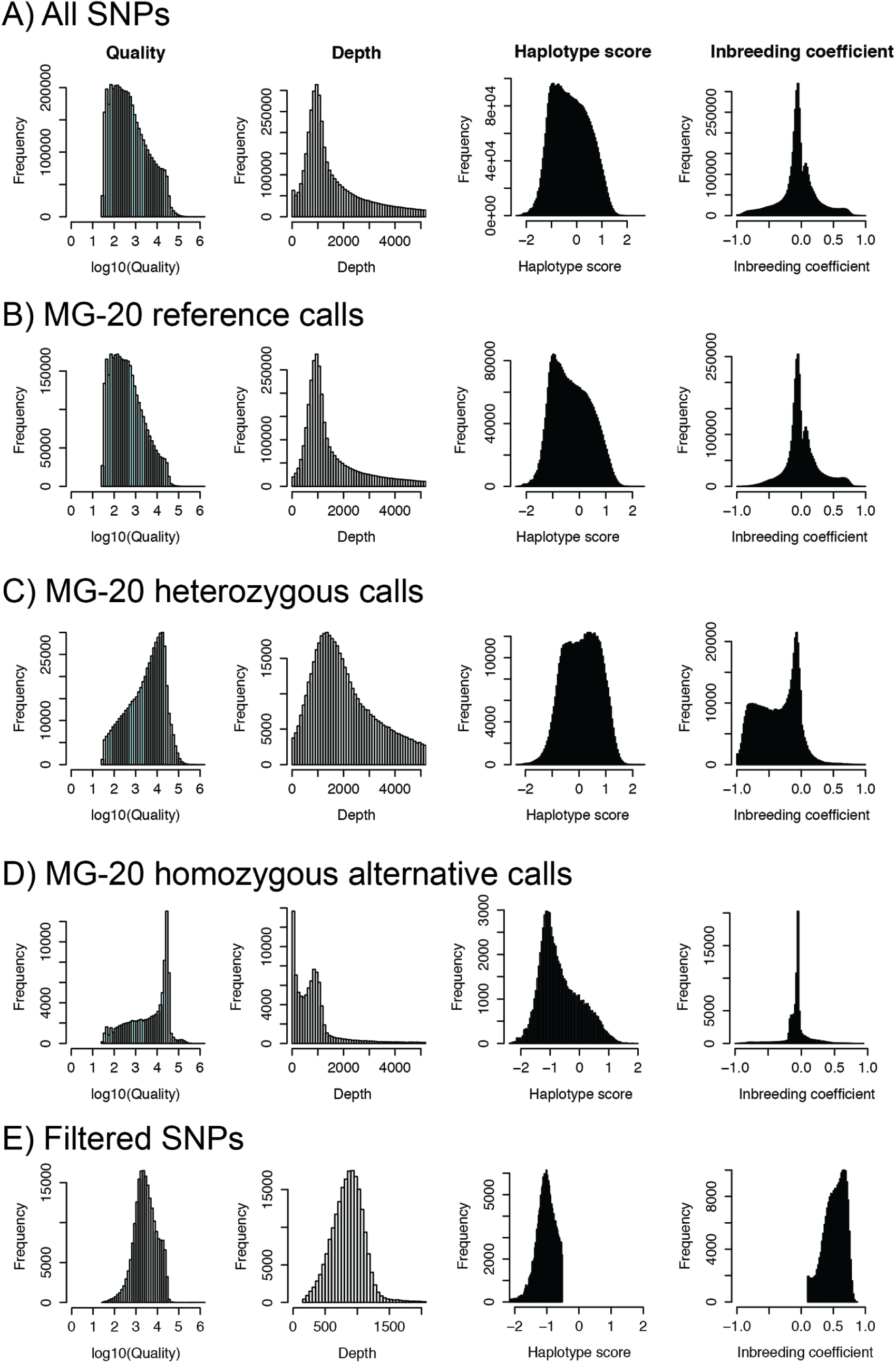
SNP property summary. Histograms for total sequencing depth, alternative allele call quality, haplotype score and inbreeding coefficients as calculated by the GATK pipeline are shown for the SNPs detected on anchored contigs. **A)** All positions. **B)** Positions with an MG20 homozygous reference call. **C)** Positions with an MG20 heterozygous call. **D)** Positions with an MG20 homozygous alternative allele call. **E)** Positions remaining after filtering requiring an MG-20 reference call, total depth > 150, Quality > 30, Haplotype score <-0.3, Inbreeding coefficient >0.1, no more than 50% missing data, and a minor allele frequency of at least 5%. The haplotype score indicates the consistency of the site with two (and only two) segregating haplotypes. Higher scores are indicative of regions with low quality alignments, which can often result in false positive SNP and indel calls. The inbreeding coefficient is the result of a likelihood-based test for the level of inbreeding.

**Supplemental figure 2:**
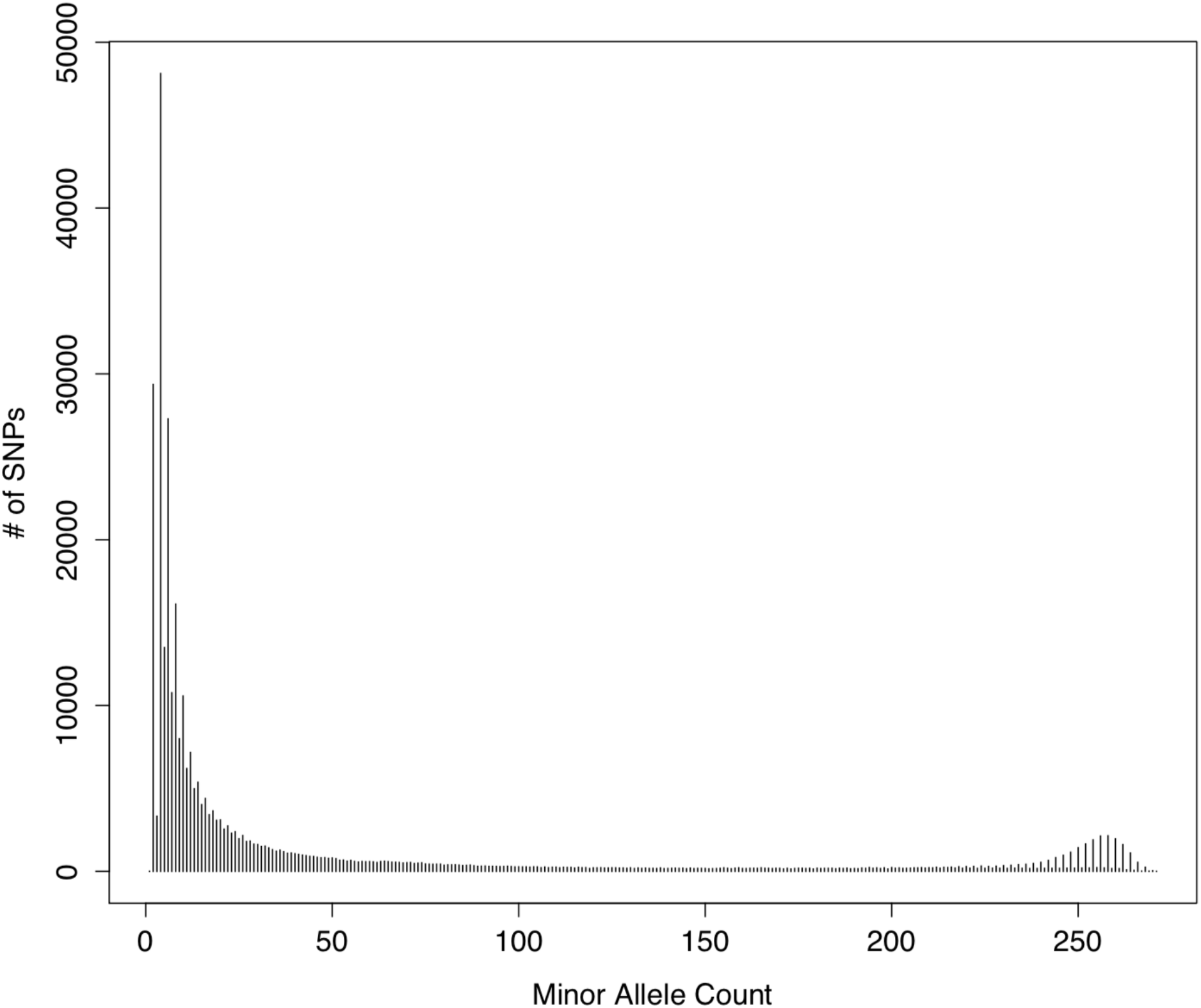
Site frequency spectrum. Non-reference allele count of the SNPs in the non-repetitive regions of the genome.

**Supplemental figure 3:**
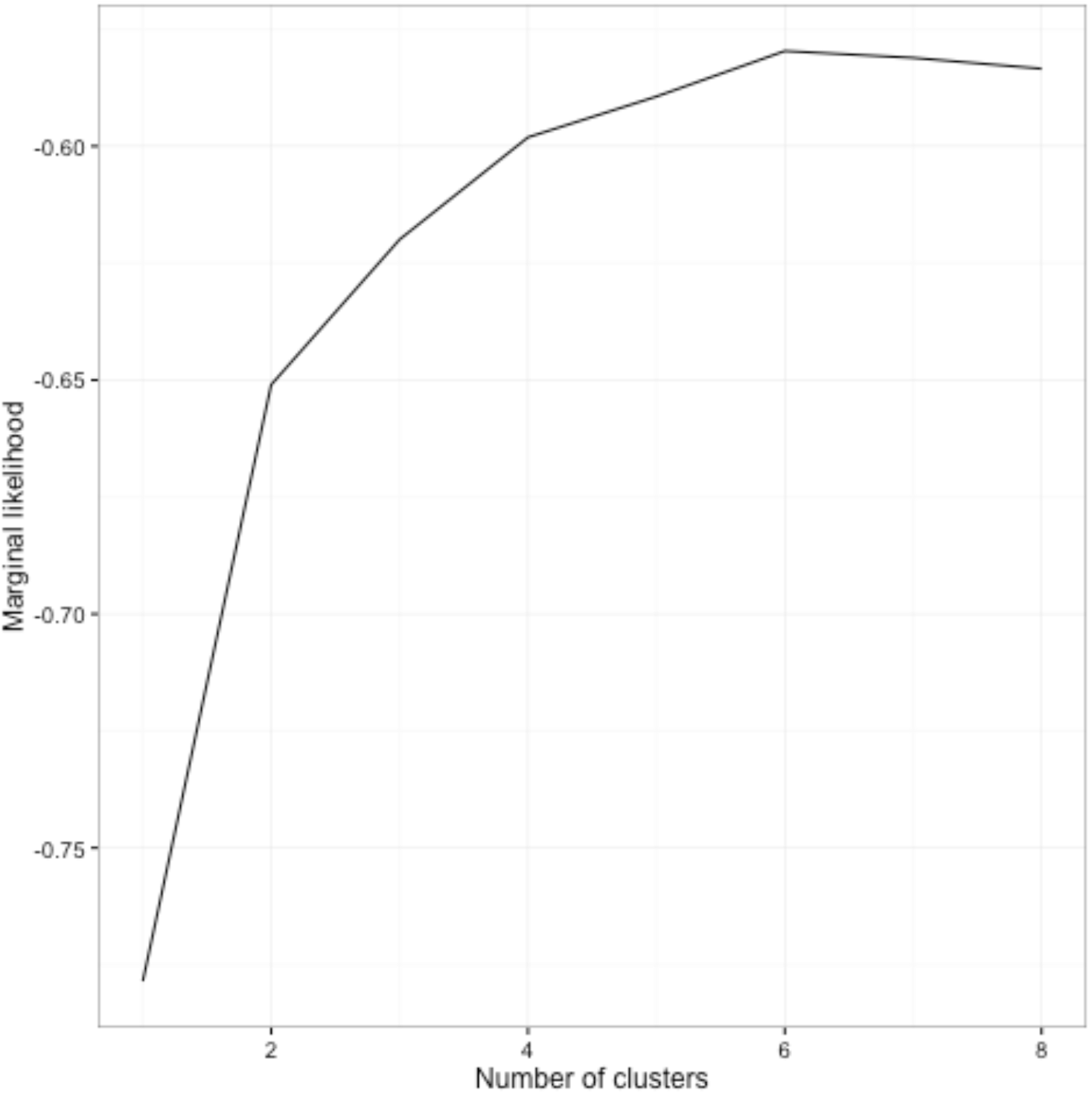
Supplemental figure 3: Estimating the number of *Lotus* subpopulations. Marginal likelihoods as a function of the number of subpopulations (*K*) in the fastSTRUCTURE analysis.

**Supplemental figure 4.**
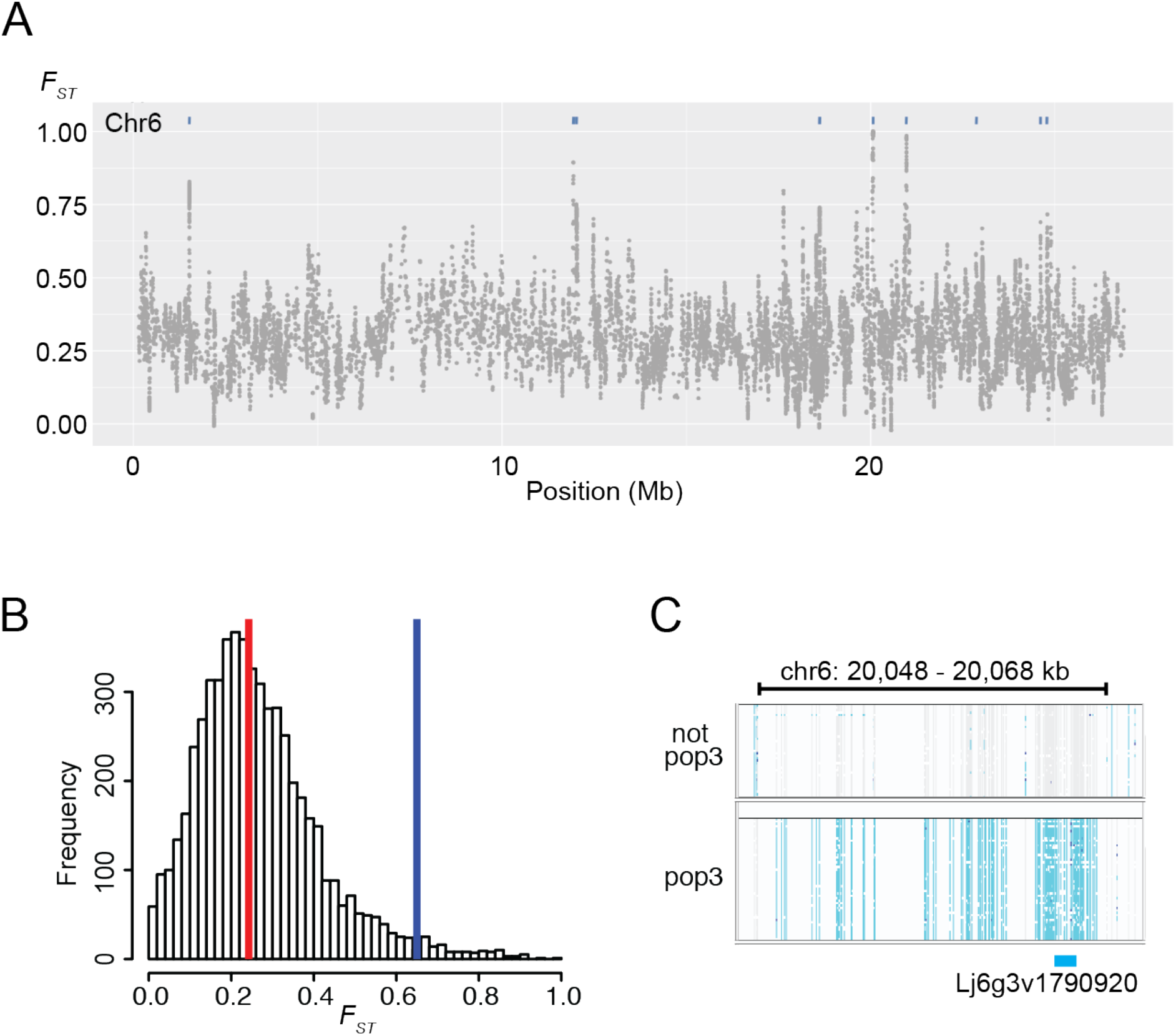
Population 3 differentiation. **A)** Fixation index (*F*_ST_) for accessions with more than 99% pop3 membership versus accessions with less than 0.1% pop3 membership. Each grey dot represents the average *F*_ST_ for 10 SNPs. Blue bars indicate the positions of genes with at least four informative SNPs and an average *F*_ST_ above 0.65. **B)** Histogram of average *F*_ST_ per gene. A total of 5,612 genes with at least 4 informative SNPs were included in the analysis. The vertical lines indicate the median *F*_ST_ (red) and an *F*_ST_ of 0.65 (blue). **C)** Genotype visualization for accessions with full or no pop3 membership in the genomic region containing the gene *Lj6g3v1790920*, which had the highest average *F*_ST_ score. Genotypes are indicated by colors: homozygous reference (grey), homozygous alternative (light blue), heterozygous (dark blue), and no call (white).

**Supplemental figure 5.**
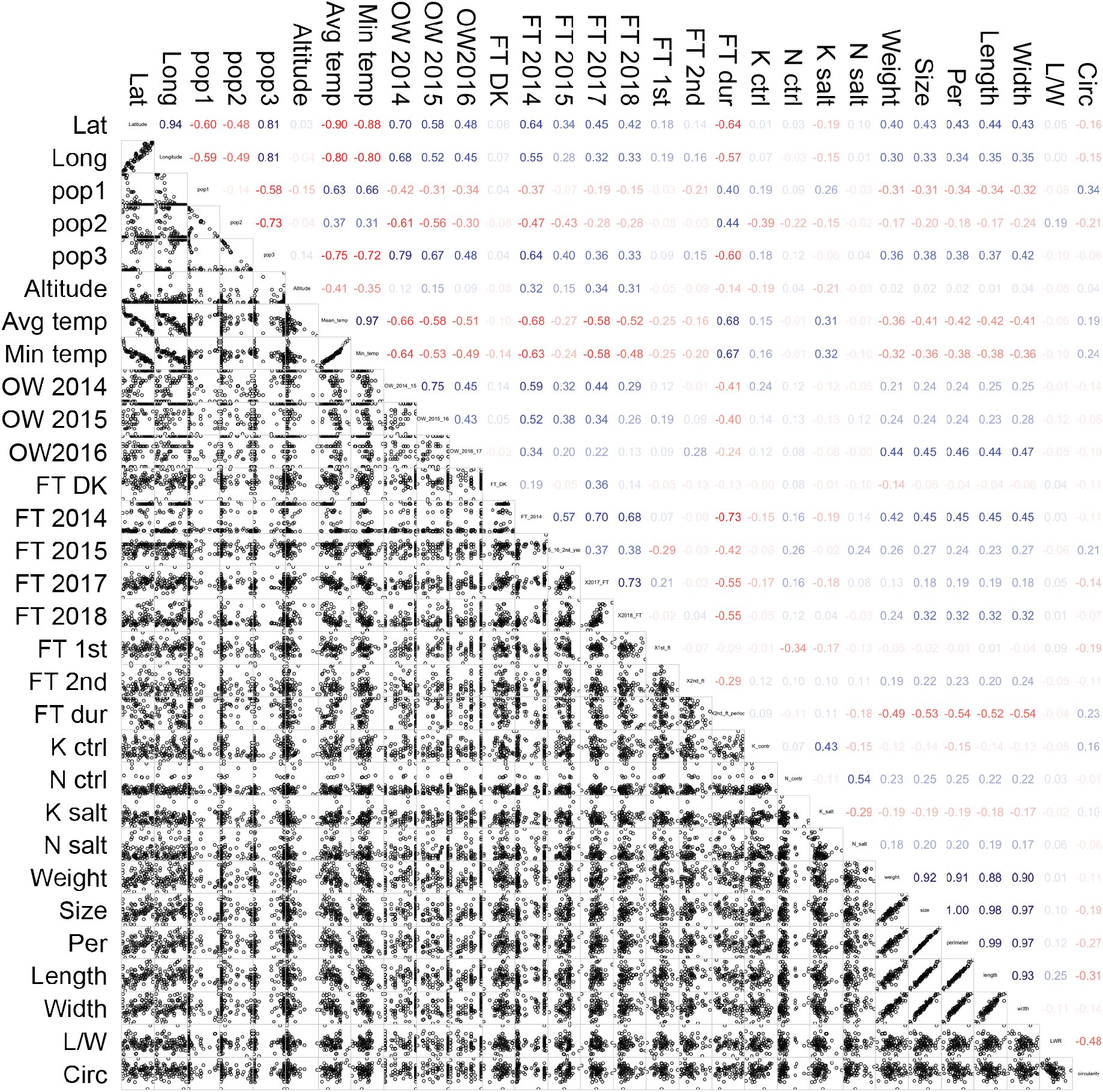
Phenotype correlations. Pairwise correlations between sets of phenotype and environmental data. Scatter plots are shown below and Pearson correlation coefficients are listed above the diagonal. Positive (blue) and negative (red) correlations are indicated. Lat: latitude. Long: longitude. pop1-3: population membership based on fastSTRUCTURE analysis (see **Figure 2** and **Supplemental file 1**). Lat, Long, Altitude, Average (Avg) and Minimum Temperature (temp) refer to accession collection sites. OW: overwintering, the fraction of plants surviving winter. FT: flowering time. FT DK: days to flowering in a greenhouse located at 56.229406N, 10.126424E, 8380, Trige, Denmark. FT 1st: weeks until first flowering period of the second year plants in 2015. FT 2nd: weeks until 2nd flowering time period of the second year plants in 2015. FT dur: duration of second flowering period of the second year plants in 2015. K^+^ and Na^+^: potassium and sodium ion accumulation in roots. ctrl: control. salt: salt treated. Weight, Size, Perimeter (Per), Length, Width and Circularity (Circ), refer to seed properties.

**Supplemental figure 6.**
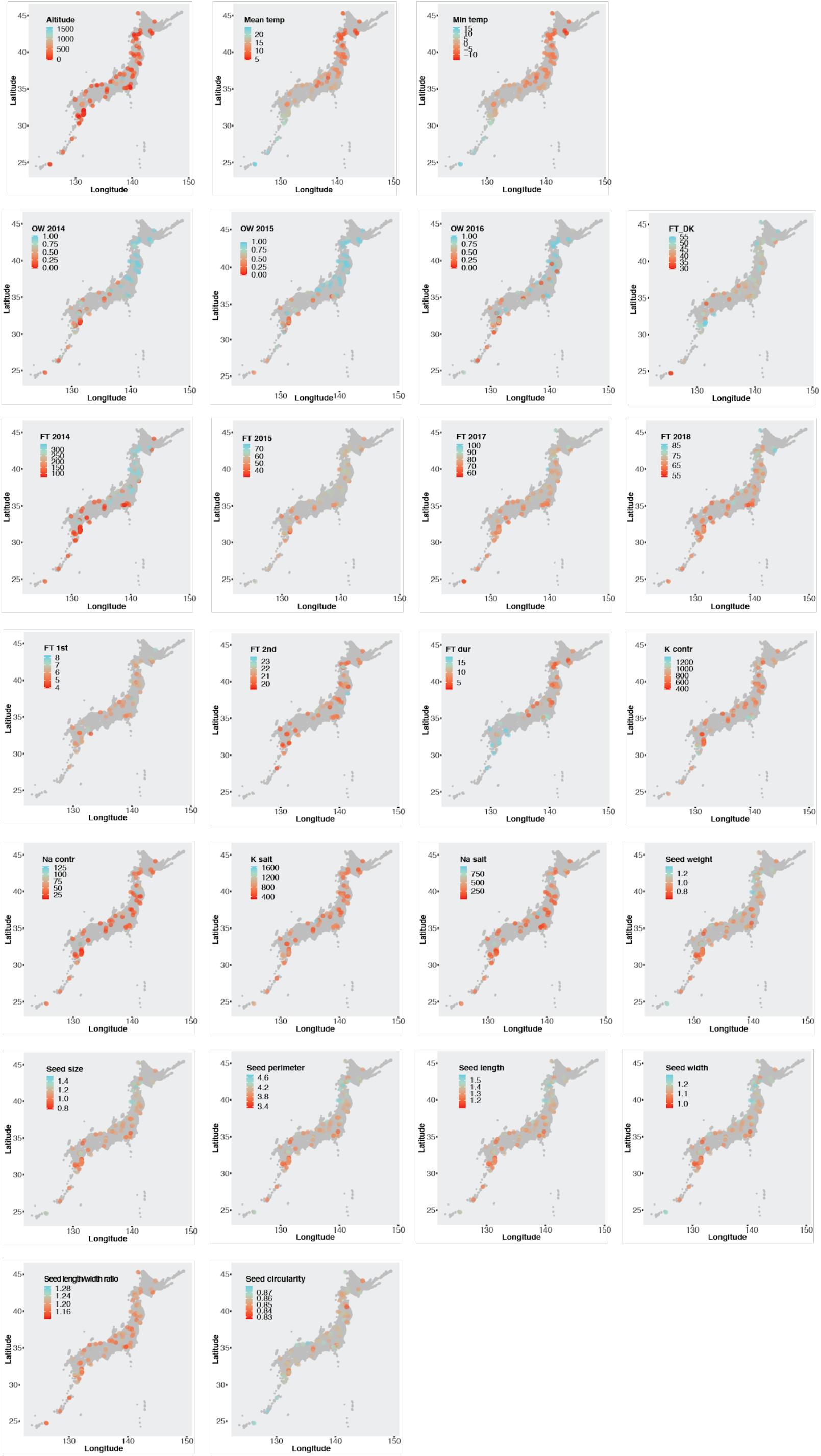
Phenotypes and geographic origin. See **Supplemental file 1** for phenotype descriptions.

**Supplemental figure 7.**
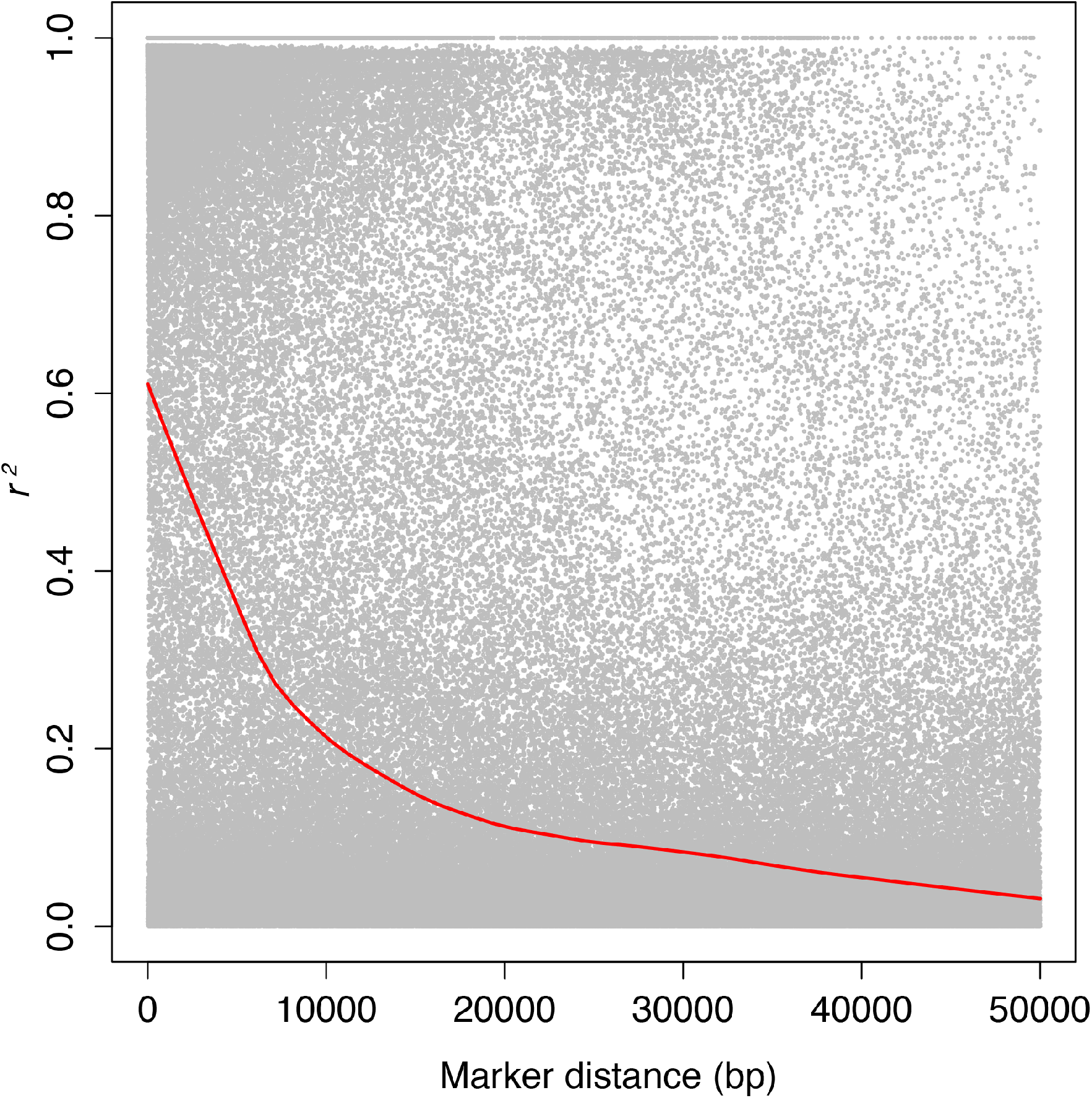
LD decay. Linkage disequilibrium (*r^2^*) as a function of pairwise marker distance.

**Supplemental figure 8.**
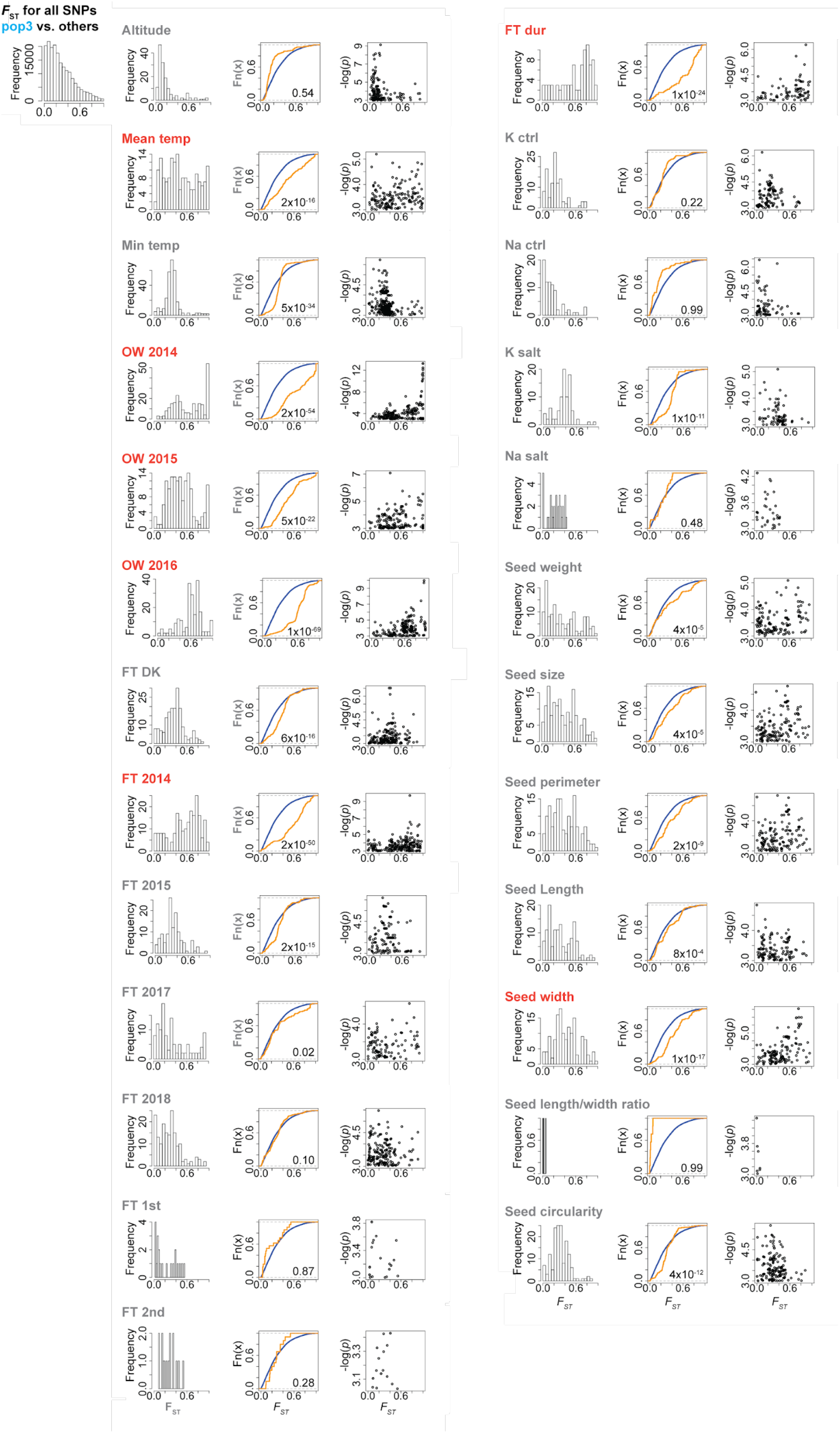
GWA and *F*_ST_ overlaps for pop3 vs non-pop3. The leftmost panel shows distributions of *F*_ST_ values for all SNPs or for the SNPs with -log(*p*) GWA scores > 3 for the traits indicated. The middle panel shows experimental cumulative distributions graphs for the *F*_ST_ values for all SNPs (blue) and the SNPs shown in the leftmost panel (orange). If the orange curve lies below the blue curve, it indicates a shift to higher *F*_ST_ values for the SNPs with high GWA scores. The rightmost panel shows *F*_ST_ values plotted versus GWA -log(*p*) scores. Altitude, Average (Avg) and Minimum (Min) temperature (temp) refer to accession collection sites. OW: overwintering, the fraction of plants surviving winter. FT: flowering time. FT DK: days to flowering in a greenhouse located at 56.229406N, 10.126424E, 8380, Trige, Denmark. FT 1st: weeks until first flowering period of the second year plants in 2015. FT 2nd: weeks until 2nd flowering time period of the second year plants in 2015. FT dur: duration of second flowering period of the second year plants in 2015. K^+^ and Na^+^: potassium and sodium ion accumulation in roots. ctrl: control. salt: salt treated. Traits likely associated with local adaptation are highlighted (orange).

**Supplemental figure 9.**
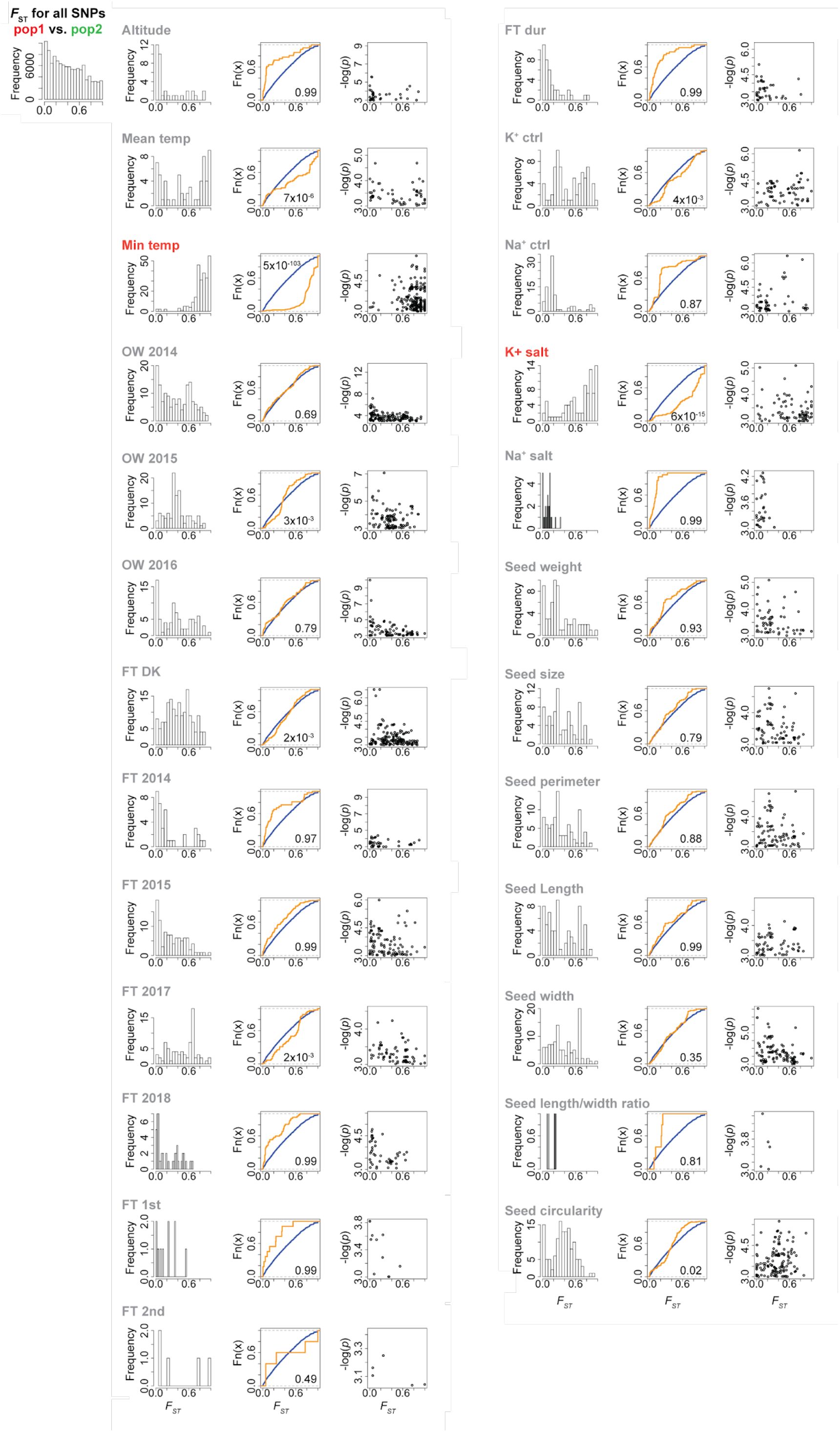
GWA and *F*_ST_ overlaps for pop1 vs pop2 comparison. The leftmost panel shows distributions of *F*_ST_ values for all SNPs (top panel) or for the SNPs with -log(*p*) GWA scores > 3 for the traits indicated. The middle panel shows experimental cumulative distributions graphs for the *F*_ST_ values for all SNPs (blue) and the SNPs shown in the leftmost panel (orange). If the orange curve lies below the blue curve, it indicates a shift to higher *F*_ST_ values for the SNPs with high GWA scores. The rightmost panel shows *F*_ST_ values plotted versus GWA -log(*p*) scores. Altitude, Average (Avg) and Minimum (Min) temperature (temp) refer to accession collection sites. OW: overwintering, the fraction of plants surviving winter. FT: flowering time. FT DK: days to flowering in a greenhouse located at 56.229406N, 10.126424E, 8380, Trige, Denmark. FT 1st: weeks until first flowering period of the second year plants in 2015. FT 2nd: weeks until 2nd flowering time period of the second year plants in 2015. FT dur: duration of second flowering period of the second year plants in 2015. K^+^ and Na^+^: potassium and sodium ion accumulation in roots. ctrl: control. salt: salt treated. Traits likely associated with local adaptation are highlighted (orange).

**Supplemental figure 10.**
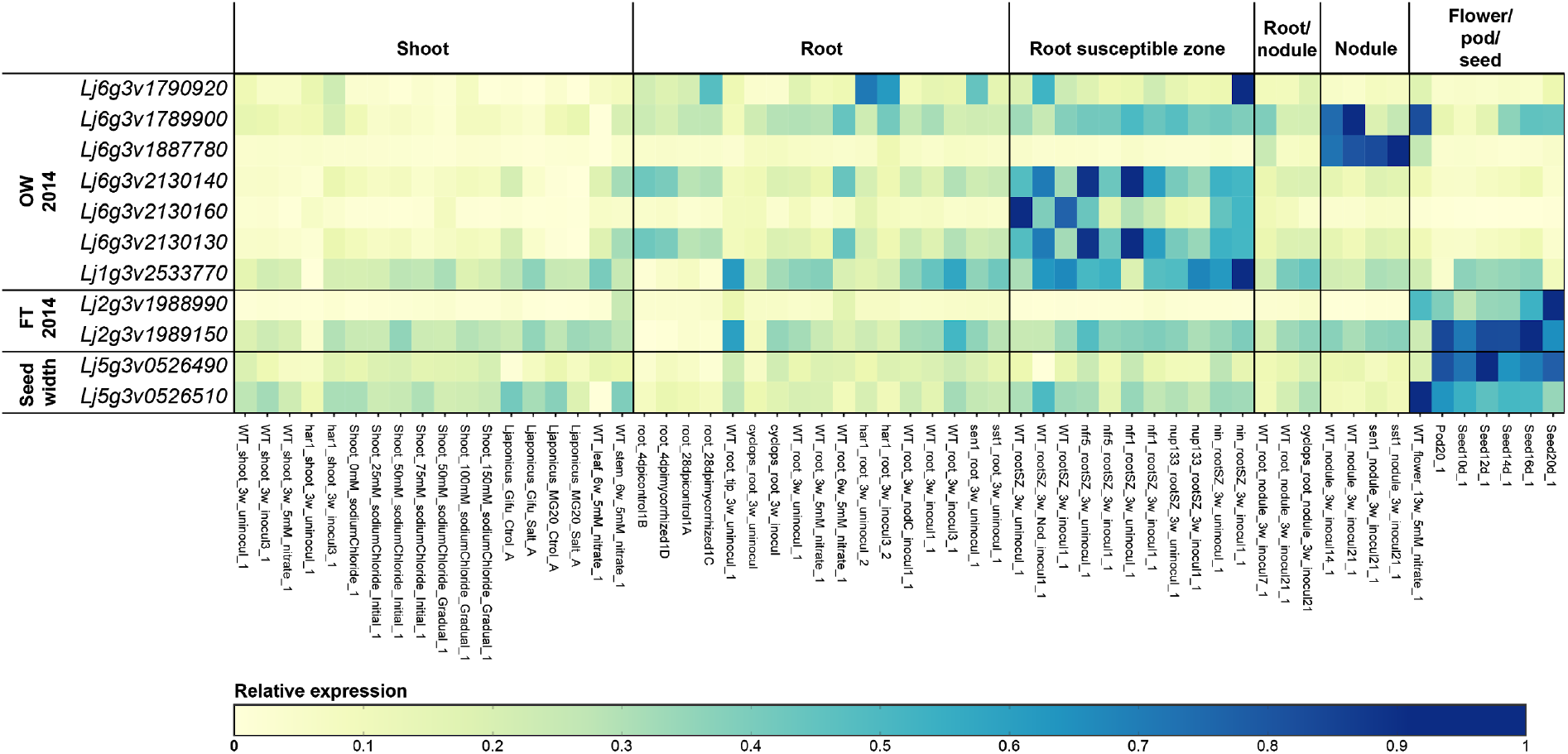
Candidate gene expression patterns. The associated trait is indicated on the left. OW: overwintering. FT: flowering time. Expression data was retrieved from Lotus Base (https://lotus.au.dk). *Lj6g3v1790920:* RING-type zinc finger protein *Lj6g3v1789900:* 1-aminocyclopropane-1-carboxylate oxidase *Lj6g3v1887780:* HBS1-like protein, Alpha/Beta hydrolase fold (IPR029058) *Lj6g3v2130140:* G-type lectin S-receptor-like serine/threonine-protein kinase *Lj6g3v2130160:* G-type lectin S-receptor-like serine/threonine-protein kinase *Lj6g3v2130130:* Glutamyl-tRNA(Gln) amidotransferase subunit A protein *Lj2g3v1988990:* glutamate dehydrogenase-like *Lj2g3v1989150: EMBRYONIC FLOWER* 2-like *Lj5g3v0526490:* mitochondrial glycoprotein family protein *Lj5g3v0526510:* putative thiol peptidase family protein

### Supplemental files

**Supplemental file 1:** Accession metadata

**Supplemental file 2:** Filtered VCF file

**Supplemental file 3:** GWA and *F*_ST_ scores for all traits

**Supplemental file 4:** GWA and *F*_ST_ overlap plots, pop3 vs. non-pop3

**Supplemental file 5:** GWA and *F*_ST_ overlap plots, pop1 vs. pop2

**Supplemental file 6:** Unix shell scripts and commands

**Supplemental file 7:** R scripts and commands

**Supplemental file 8:** Imputed genotype file used for GWA analysis

