## Supplementary material for "Extreme genetic signatures of local adaptation during plant colonization"

**Supplemental file 4:** GWA and $F_{ST}$ (pop3 vs. non-pop3) overlaps for all traits and chromosomes $F_{ST}$ averages for 10 SNPs are indicated by grey dots. In the left panel, colored dots indicate $F_{ST}$ values for SNPs with GWA -log(*p*) scores > 3 for the chromosomes and traits indicated. In the right panel $F_{ST}$ averages for 10 SNPs (grey dots) are overlaid with GWAS -log(*p*) scores > 3 for the chromosomes and traits indicated. Each chromosome is shown on a separate page. Viewing at 500% is recommended.

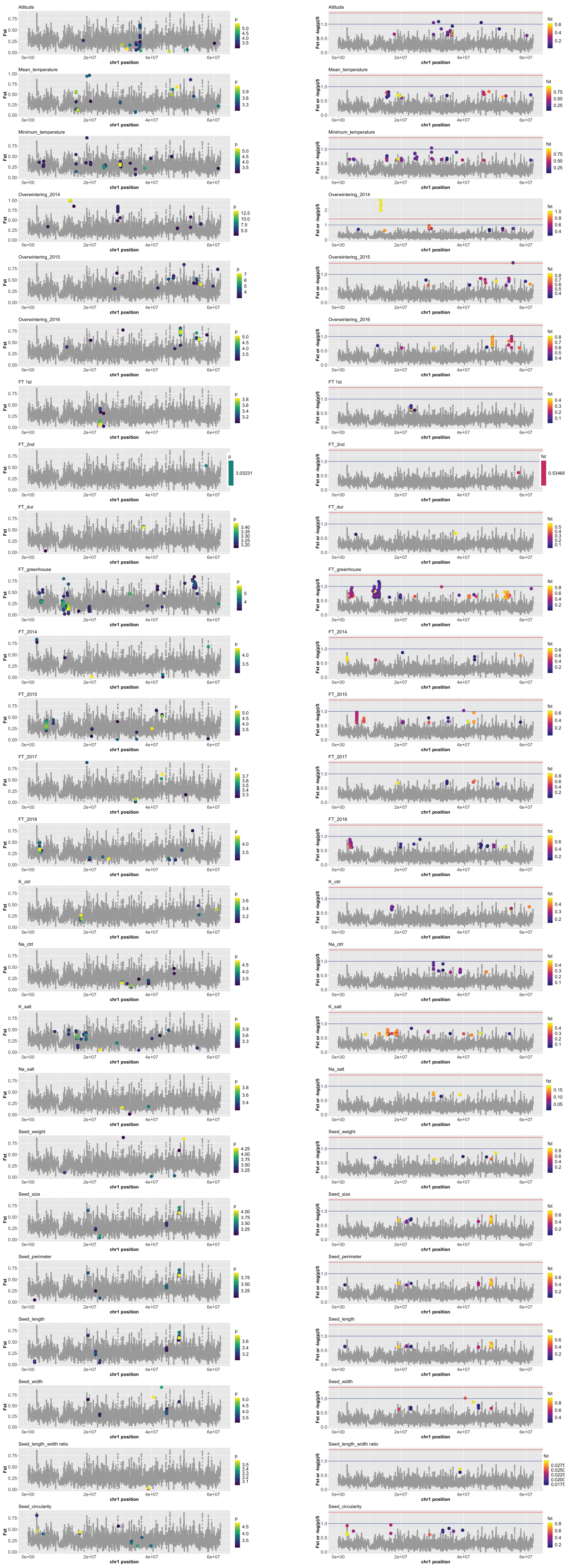
Altitude
Mean_temperature
Minimum_temperature
Overwintering_2014
Overwintering_2015
Overwintering_2016
FT 1st
FT_2nd
FT_dur
FT_greenhouse
FT_2014
FT_2015
FT_2017
FT_2018
K_ctrl
Na_ctrl
K_salt
Na_salt
Seed_weight
Seed_size
Seed_perimeter
Seed_length
Seed_width
Seed_length_width ratio
Seed_circularity
Fst
Fst or -log(p)/5
chr1 position
p
fst

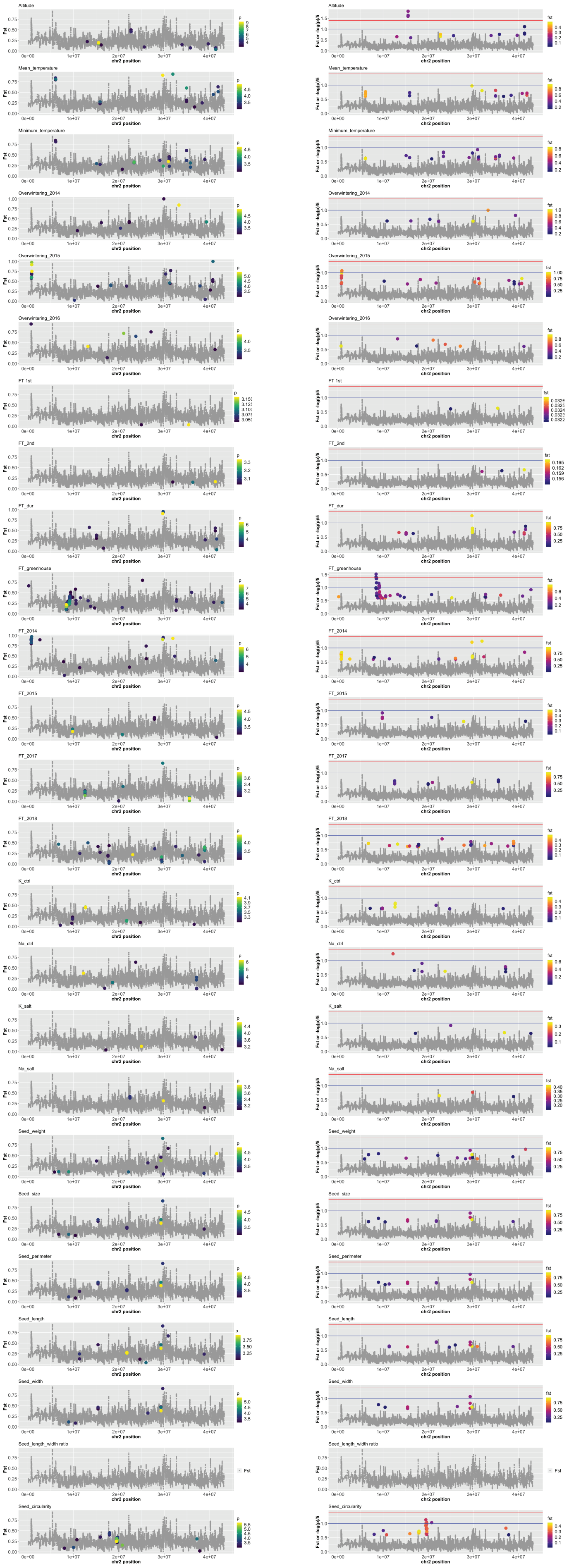
Altitude
Mean_temperature
Minimum_temperature
Overwintering_2014
Overwintering_2015
Overwintering_2016
FT 1st
FT_2nd
FT_dur
FT_greenhouse
FT_2014
FT_2015
FT_2017
FT_2018
K_ctrl
Na_ctrl
K_salt
Na_salt
Seed_weight
Seed_size
Seed_perimeter
Seed_length
Seed_width
Seed_length_width ratio
Seed_circularity
Fst
Fst or -log(p)/5
chr2 position
p
fst

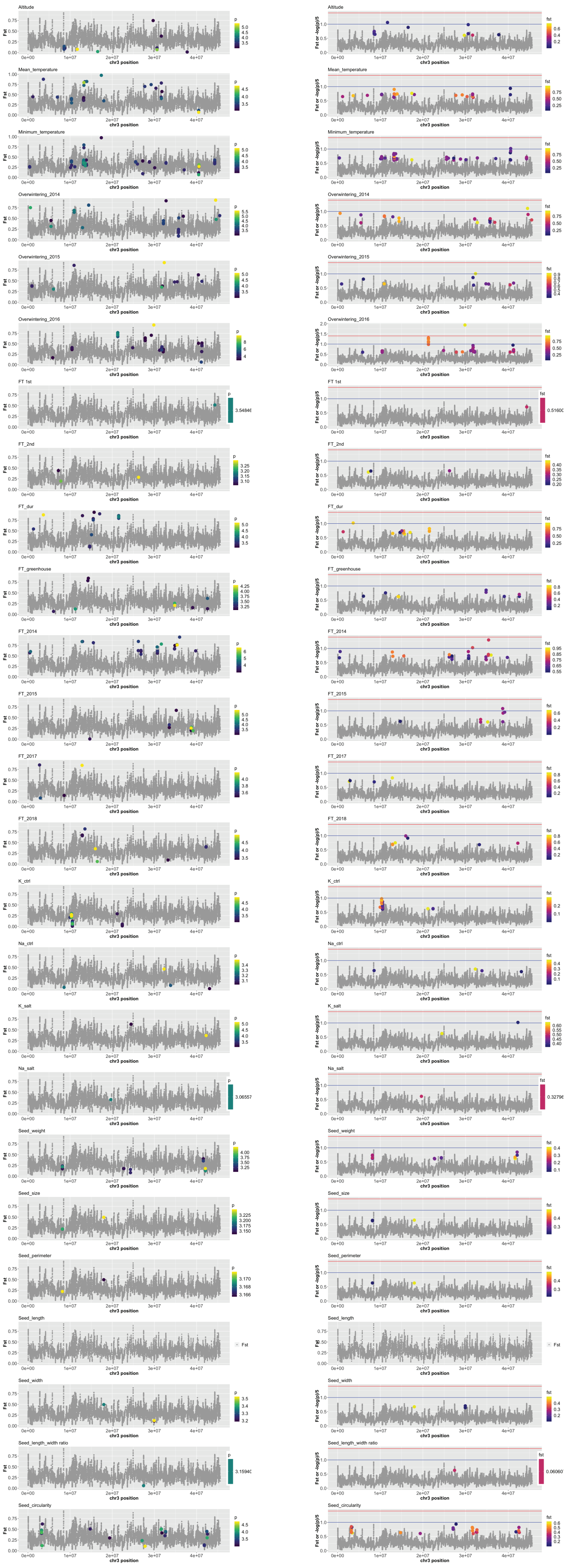

Altitude
Mean_temperature
Minimum_temperature
Overwintering_2014
Overwintering_2015
Overwintering_2016
FT 1st
FT_2nd
FT_dur
FT_greenhouse
FT_2014
FT_2015
FT_2017
FT_2018
K_ctrl
Na_ctrl
K_salt
Na_salt
Seed_weight
Seed_size
Seed_perimeter
Seed_length
Seed_width
Seed_length_width ratio
Seed_circularity
Fst
Fst or -log(p)/5
chr3 position

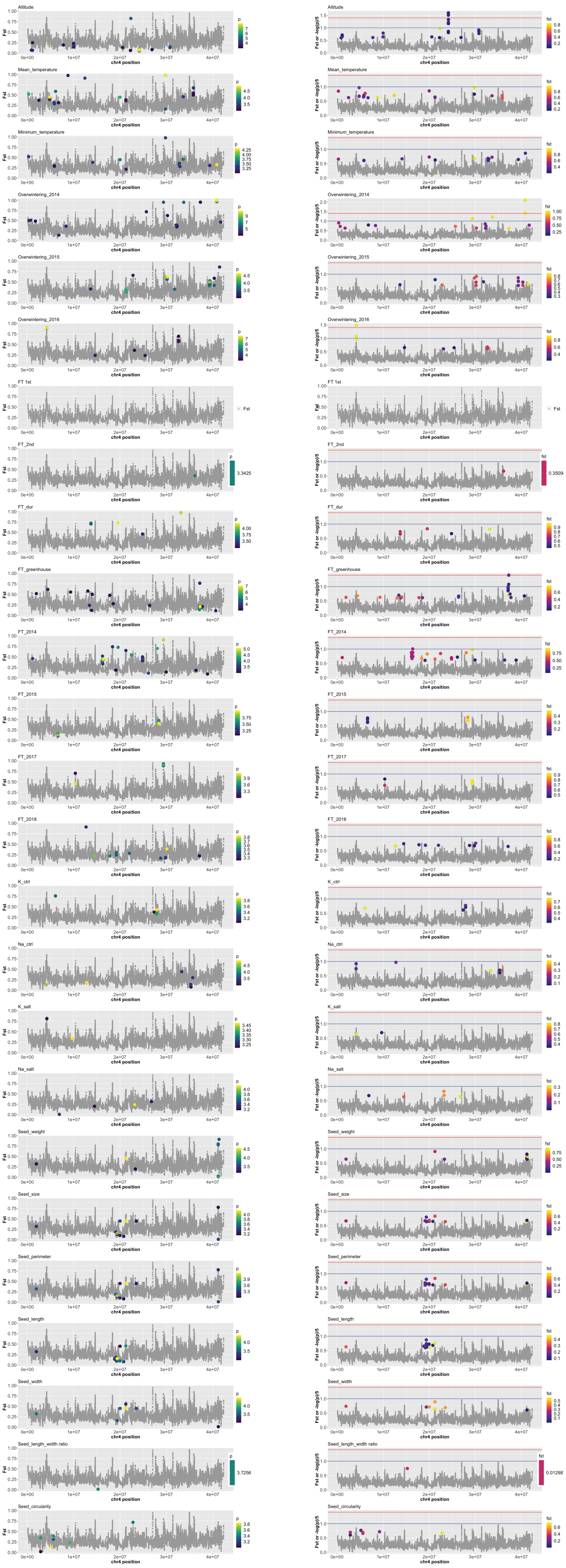

Altitude
Mean_temperature
Minimum_temperature
Overwintering_2014
Overwintering_2015
Overwintering_2016
FT 1st
FT_2nd
FT_dur
FT_greenhouse
FT_2014
FT_2015
FT_2017
FT_2018
K_ctrl
Na_ctrl
K_salt
Na_salt
Seed_weight
Seed_size
Seed_perimeter
Seed_length
Seed_width
Seed_length_width ratio
Seed_circularity
Fst
Fst or -log(p)/5
chr4 position
p
fst

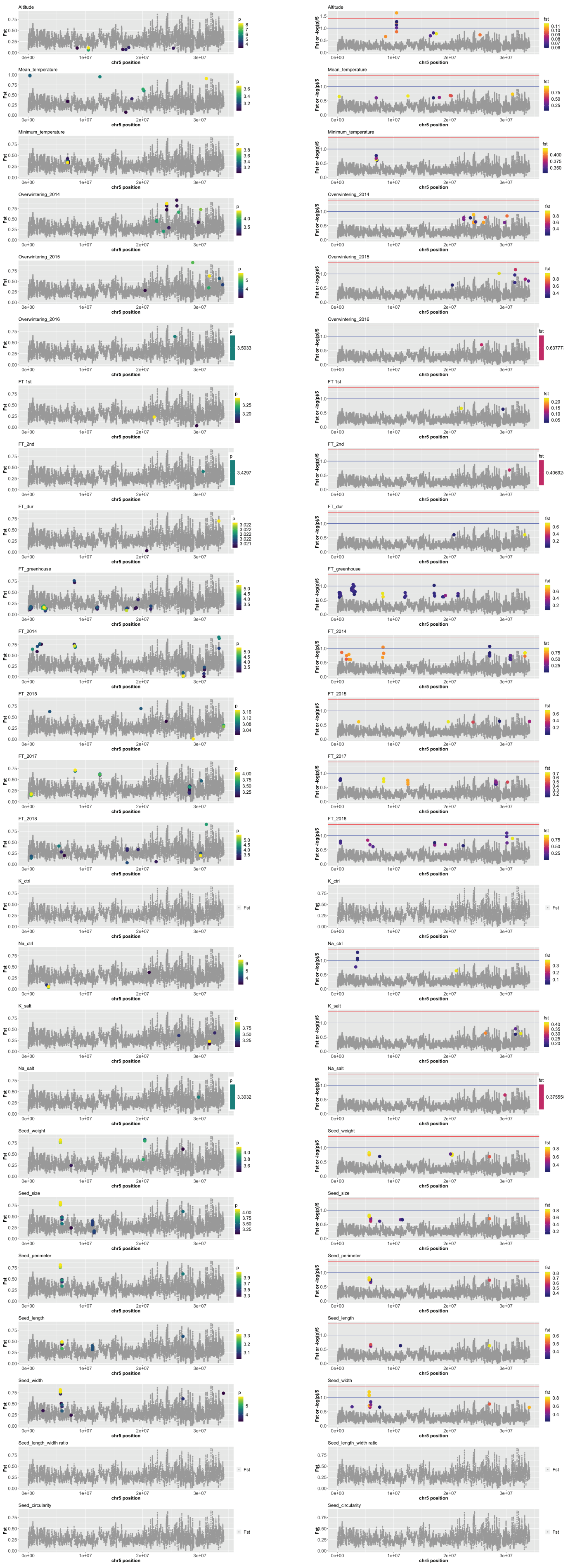
Altitude
Mean_temperature
Minimum_temperature
Overwintering_2014
Overwintering_2015
Overwintering_2016
FT 1st
FT_2nd
FT_dur
FT_greenhouse
FT_2014
FT_2015
FT_2017
FT_2018
K_ctrl
Na_ctrl
K_salt
Na_salt
Seed_weight
Seed_size
Seed_perimeter
Seed_length
Seed_width
Seed_length_width ratio
Seed_circularity
Fst
Fst or -log(p)/5
chr5 position
p
fst

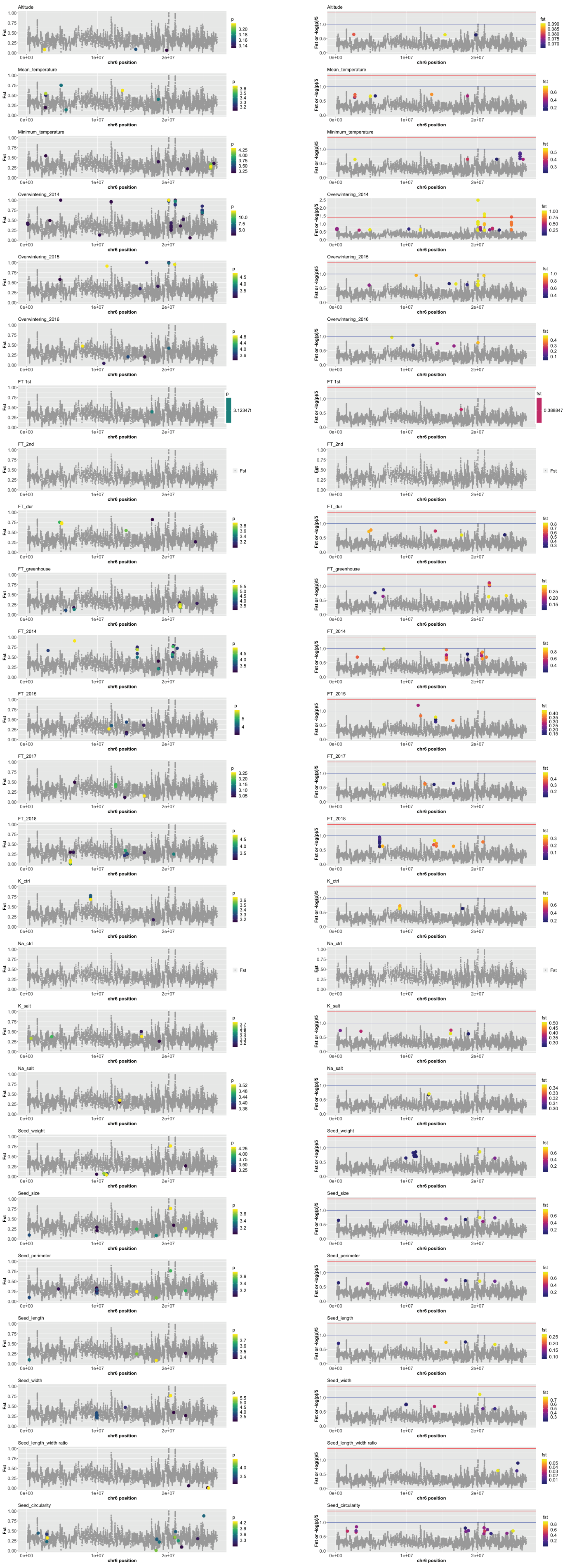
Altitude
Mean_temperature
Minimum_temperature
Overwintering_2014
Overwintering_2015
Overwintering_2016
FT 1st
FT_2nd
FT_dur
FT_greenhouse
FT_2014
FT_2015
FT_2017
FT_2018
K_ctrl
Na_ctrl
K_salt
Na_salt
Seed_weight
Seed_size
Seed_perimeter
Seed_length
Seed_width
Seed_length_width ratio
Seed_circularity
Fst
Fst or -log(p)/5
chr6 position
p
fst
