## Supplementary material for "Extreme genetic signatures of local adaptation during plant colonization"

### ----------Variant calling----------------###
{
################################## Mapping ##################################
bwa mem -t 15 lj_r30.fa MG-147_S8_L001_R1_001.fastq MG-147_S8_L001_R2_001.fastq > MG-147_L001.sam


################################## Sam To Bam ##################################
samtools view -bS MG-147_L001.sam | samtools sort - MG-147_L001_sorted


################################## Add readgroup ##################################
java -jar /com/extra/picard/1.96/jar-bin/AddOrReplaceReadGroups.jar I=MG-147.bam O=MG-147.rg.bam SO=coordinate ID=MG-147 LB=MG-147 PL=MiSeq PU=one SM=MG-147


################################## Find coverage from bam file ##################################
samtools depth -a $bamfile | awk '{sum+=$3; } END { print "Average = ",sum/NR; }'


################################## Picard and GATK steps ##################################
#1 Mark duplicates
qx --no-scratch -n=1 -c=1 -m =26g -t=50:00:00 'java -Xmx26g -jar /com/extra/picard/2.7.1/jar-lib/picard.jar MarkDuplicates INPUT=Gifu.bam OUTPUT=Gifu.dedup.bam METRICS_FILE=Gifu.metrics.txt'

#2 index dedup file
qx --no-scratch -n=1 -c=1 -t=50:00:00 '/com/extra/samtools/0.1.19/bin/samtools index Gifu.dedup.bam Gifu.dedup.bai'

#3 ReAlign
qx --no-scratch -n=1 -c=1 -m=20g -t=24:00:00 'gatk -U -T RealignerTargetCreator -R /home/niraj/LotusGenome/100_data/124Accessions/BAMs/ljFiles/lj_r30.fa -I Gifu.dedup.bam -o Gifu.intervals'

#4 IndelRealign
qx --no-scratch -n=1 -c=1 -m=25g -t=24:00:00 'gatk -U -T IndelRealigner -R /home/niraj/LotusGenome/100_data/124Accessions/BAMs/ljFiles/lj_r30.fa -targetIntervals Gifu.intervals -I Gifu.dedup.bam -o Gifu.realigned.bam'

#5 UnifiedGenotyper  (This can be run right in the beginning)
qx --no-scratch -n=1 -c=1 -m=20g -t=100:00:00 'gatk -T UnifiedGenotyper -nt 1 -glm BOTH -R /home/niraj/LotusGenome/100_data/124Accessions/BAMs/ljFiles/lj_r30.fa -I Gifu.bam -I MG001.bam -I MG002.bam -I MG003.bam -I MG004.bam -I MG005.bam -I MG007.bam -I MG008.bam -I MG009.bam -I MG010.bam  -I MG011.bam -I MG012.bam -I MG013.bam -I MG014.bam -I MG016.bam -I MG017.bam -I MG018.bam  -I MG019.bam  -I MG020.bam -I MG021.bam  -I MG022.bam  -I MG023.bam -I MG024.bam  -I MG025.bam -I MG026.bam  -I MG027.bam -I MG028.bam  -I MG030.bam  -I MG032.bam  -I MG033.bam  -I MG034.bam -I MG035.bam  -I MG036.bam  -I MG038.bam  -I MG039.bam  -I MG040.bam -I MG041.bam -I MG042.bam -I MG044.bam -I MG045.bam -I MG046.bam -I MG049.bam -I MG050.bam -I MG051.bam -I MG052.bam -I MG053.bam -I MG056.bam -I MG057.bam -I MG058.bam -I MG059.bam -I MG060.bam -I MG061.bam -I MG062.bam -I MG063.bam -I MG065.bam -I MG066.bam -I MG067.bam -I MG068.bam -I MG069.bam -I MG070.bam -I MG071.bam -I MG072.bam -I MG073.bam -I MG074.bam -I MG075.bam -I MG076.bam -I MG077.bam -I MG078.bam -I MG080.bam -I MG081.bam -I MG082.bam -I MG083.bam -I MG084.bam -I MG085.bam -I MG086.bam -I MG088.bam -I MG089.bam -I MG090.bam -I MG091.bam -I MG092.bam -I MG093.bam -I MG094.bam -I MG095.bam -I MG096.bam -I MG097.bam -I MG098.bam -I MG099.bam -I MG100.bam -I MG101.bam -I MG102.bam -I MG103.bam -I MG104.bam -I MG105.bam -I MG106.bam -I MG107.bam -I MG109.bam -I MG110.bam -I MG111.bam -I MG112.bam  -I MG113.bam -I MG115.bam -I MG116.bam -I MG117.bam -I MG118.bam -I MG119.bam -I MG120.bam -I MG121.bam -I MG123.bam -I MG124.bam -I MG125.bam -I MG126.bam -I MG127.bam -I MG128.bam -I MG129.bam  -I MG130.bam -I MG131.bam -I MG132.bam -I MG133.bam -I MG134.bam -I MG135.bam -I MG136.bam -I MG138.bam -I MG139.bam -I MG140.bam -I MG141.bam -I MG142.bam -I MG143.bam -I MG144.bam -I MG145.bam -I MG146.bam -I MG149.bam -I MG150.bam -I MG151.bam -I MG152.bam -I MG154.bam -I MG155.bam -I MG156.bam -o 137Lj.vcf -stand_call_conf 90 -dcov 200' 

#6 BaseRecalibrator
qx --no-scratch -n=1 -c=1 -m=15g -t=50:00:00 'gatk -T BaseRecalibrator -rf BadCigar -knownSites 137LjAccessions.vcf -R /home/niraj/LotusGenome/100_data/124Accessions/BAMs/ljFiles/lj_r30.fa -I Gifu.realigned.bam -o Gifu.recal.table’

#7 Print Reads
qx --no-scratch -n=1 -c=1 -m=20g -t=24:00:00 'java -Djava.awt.headless=true -Xmx8g -jar /com/extra/GATK/2.7-2/jar-bin/GenomeAnalysisTK.jar -et NO_ET -K /com/extra/GATK/2.7-2/runef_birc.au.dk.key -T PrintReads -R /home/niraj/LotusGenome/100_data/124Accessions/BAMs/ljFiles/lj_r30.fa -I Gifu.realigned.bam -BQSR Gifu.recal.table -o Gifu.recalreads.bam'

#8 ReduceReads
qx --no-scratch -n=1 -c=1 -m=20g -t=24:00:00 'java -Djava.awt.headless=true -Djava.io.tmpdir=/home/niraj/LotusGenome/Niraj/ -Xmx6g -jar /com/extra/GATK/2.7-2/jar-bin/GenomeAnalysisTK.jar -et NO_ET -K /com/extra/GATK/2.7-2/runef_birc.au.dk.key -T ReduceReads -rf BadCigar -R /home/niraj/LotusGenome/100_data/124Accessions/BAMs/ljFiles/lj_r30.fa -I Gifu.recalreads.bam -o Gifu.reduced.bam'

#9 Re unifiedgenotyper
qx --no-scratch -n=1 -c=5 -m=20g -t=100:00:00 'java -Djava.awt.headless=true -Djava.io.tmpdir=/home/niraj/LotusGenome/Niraj/ -Xmx12g -jar /com/extra/GATK/2.7-2/jar-bin/GenomeAnalysisTK.jar -et NO_ET -K /com/extra/GATK/2.7-2/runef_birc.au.dk.key -T UnifiedGenotyper -nt 5 -L chr1 -glm BOTH -R /home/niraj/LotusGenome/100_data/124Accessions/BAMs/ljFiles/lj_r30.fa -I Gifu.reduced.bam -I MG001.reduced.bam -I MG002.reduced.bam -I MG003.reduced.bam -I MG004.reduced.bam -I MG005.reduced.bam -I MG007.reduced.bam -I MG008.reduced.bam -I MG009.reduced.bam -I MG010.reduced.bam  -I MG011.reduced.bam -I MG012.reduced.bam -I MG013.reduced.bam -I MG014.reduced.bam -I MG016.reduced.bam -I MG017.reduced.bam -I MG018.reduced.bam  -I MG019.reduced.bam  -I MG020.reduced.bam -I MG021.reduced.bam  -I MG022.reduced.bam  -I MG023.reduced.bam -I MG024.reduced.bam  -I MG025.reduced.bam -I MG026.reduced.bam  -I MG027.reduced.bam -I MG028.reduced.bam  -I MG030.reduced.bam  -I MG032.reduced.bam  -I MG033.reduced.bam  -I MG034.reduced.bam -I MG035.reduced.bam  -I MG036.reduced.bam  -I MG038.reduced.bam  -I MG039.reduced.bam  -I MG040.reduced.bam -I MG041.reduced.bam -I MG042.reduced.bam -I MG044.reduced.bam -I MG045.reduced.bam -I MG046.reduced.bam -I MG049.reduced.bam -I MG050.reduced.bam -I MG051.reduced.bam -I MG052.reduced.bam -I MG053.reduced.bam -I MG056.reduced.bam -I MG057.reduced.bam -I MG058.reduced.bam -I MG059.reduced.bam -I MG060.reduced.bam -I MG061.reduced.bam -I MG062.reduced.bam -I MG063.reduced.bam -I MG065.reduced.bam -I MG066.reduced.bam -I MG067.reduced.bam -I MG068.reduced.bam -I MG069.reduced.bam -I MG070.reduced.bam -I MG071.reduced.bam -I MG072.reduced.bam -I MG073.reduced.bam -I MG074.reduced.bam -I MG075.reduced.bam -I MG076.reduced.bam -I MG077.reduced.bam -I MG078.reduced.bam -I MG080.reduced.bam -I MG081.reduced.bam -I MG082.reduced.bam -I MG083.reduced.bam -I MG084.reduced.bam -I MG085.reduced.bam -I MG086.reduced.bam -I MG088.reduced.bam -I MG089.reduced.bam -I MG090.reduced.bam -I MG091.reduced.bam -I MG092.reduced.bam -I MG093.reduced.bam -I MG094.reduced.bam -I MG095.reduced.bam -I MG096.reduced.bam -I MG097.reduced.bam -I MG098.reduced.bam -I MG099.reduced.bam -I MG100.reduced.bam -I MG101.reduced.bam -I MG102.reduced.bam -I MG103.reduced.bam -I MG104.reduced.bam -I MG105.reduced.bam -I MG106.reduced.bam -I MG107.reduced.bam -I MG109.reduced.bam -I MG110.reduced.bam -I MG111.reduced.bam -I MG112.reduced.bam  -I MG113.reduced.bam -I MG115.reduced.bam -I MG116.reduced.bam -I MG117.reduced.bam -I MG118.reduced.bam -I MG119.reduced.bam -I MG120.reduced.bam -I MG121.reduced.bam -I MG123.reduced.bam -I MG124.reduced.bam -I MG125.reduced.bam -I MG126.reduced.bam -I MG127.reduced.bam -I MG128.reduced.bam -I MG129.reduced.bam  -I MG130.reduced.bam -I MG131.reduced.bam -I MG132.reduced.bam -I MG133.reduced.bam -I MG134.reduced.bam -I MG135.reduced.bam -I MG136.reduced.bam -I MG138.reduced.bam -I MG139.reduced.bam -I MG140.reduced.bam -I MG141.reduced.bam -I MG142.reduced.bam -I MG143.reduced.bam -I MG144.reduced.bam -I MG145.reduced.bam -I MG146.reduced.bam -I MG149.reduced.bam -I MG150.reduced.bam -I MG151.reduced.bam -I MG152.reduced.bam -I MG154.reduced.bam -I MG155.reduced.bam -I MG156.reduced.bam -o FinalGeno_137Lj_chr1.vcf'

## command to obtain genotypes at all the sites to calculate callable fraction
qx --no-scratch -n=1 -c=5 -m=20g -t=200:00:00 'java -Djava.awt.headless=true -Djava.io.tmpdir=/home/niraj/LotusGenome/Niraj/ -Xmx12g -jar /com/extra/GATK/2.7-2/jar-bin/GenomeAnalysisTK.jar -et NO_ET -K /com/extra/GATK/2.7-2/runef_birc.au.dk.key -T UnifiedGenotyper -nt 5 -glm BOTH -R /home/niraj/LotusGenome/100_data/124Accessions/BAMs/ljFiles/lj_r30.fa -I Gifu.reduced.bam -I MG001.reduced.bam -I MG002.reduced.bam -I MG003.reduced.bam -I MG004.reduced.bam -I MG005.reduced.bam -I MG007.reduced.bam -I MG008.reduced.bam -I MG009.reduced.bam -I MG010.reduced.bam  -I MG011.reduced.bam -I MG012.reduced.bam -I MG013.reduced.bam -I MG014.reduced.bam -I MG016.reduced.bam -I MG017.reduced.bam -I MG018.reduced.bam  -I MG019.reduced.bam  -I MG020.reduced.bam -I MG021.reduced.bam  -I MG022.reduced.bam  -I MG023.reduced.bam -I MG024.reduced.bam  -I MG025.reduced.bam -I MG026.reduced.bam  -I MG027.reduced.bam -I MG028.reduced.bam  -I MG030.reduced.bam  -I MG032.reduced.bam  -I MG033.reduced.bam  -I MG034.reduced.bam -I MG035.reduced.bam  -I MG036.reduced.bam  -I MG038.reduced.bam  -I MG039.reduced.bam  -I MG040.reduced.bam -I MG041.reduced.bam -I MG042.reduced.bam -I MG044.reduced.bam -I MG045.reduced.bam -I MG046.reduced.bam -I MG049.reduced.bam -I MG050.reduced.bam -I MG051.reduced.bam -I MG052.reduced.bam -I MG053.reduced.bam -I MG056.reduced.bam -I MG057.reduced.bam -I MG058.reduced.bam -I MG059.reduced.bam -I MG060.reduced.bam -I MG061.reduced.bam -I MG062.reduced.bam -I MG063.reduced.bam -I MG065.reduced.bam -I MG066.reduced.bam -I MG067.reduced.bam -I MG068.reduced.bam -I MG069.reduced.bam -I MG070.reduced.bam -I MG071.reduced.bam -I MG072.reduced.bam -I MG073.reduced.bam -I MG074.reduced.bam -I MG075.reduced.bam -I MG076.reduced.bam -I MG077.reduced.bam -I MG078.reduced.bam -I MG080.reduced.bam -I MG081.reduced.bam -I MG082.reduced.bam -I MG083.reduced.bam -I MG084.reduced.bam -I MG085.reduced.bam -I MG086.reduced.bam -I MG088.reduced.bam -I MG089.reduced.bam -I MG090.reduced.bam -I MG091.reduced.bam -I MG092.reduced.bam -I MG093.reduced.bam -I MG094.reduced.bam -I MG095.reduced.bam -I MG096.reduced.bam -I MG097.reduced.bam -I MG098.reduced.bam -I MG099.reduced.bam -I MG100.reduced.bam -I MG101.reduced.bam -I MG102.reduced.bam -I MG103.reduced.bam -I MG104.reduced.bam -I MG105.reduced.bam -I MG106.reduced.bam -I MG107.reduced.bam -I MG109.reduced.bam -I MG110.reduced.bam -I MG111.reduced.bam -I MG112.reduced.bam  -I MG113.reduced.bam -I MG115.reduced.bam -I MG116.reduced.bam -I MG117.reduced.bam -I MG118.reduced.bam -I MG119.reduced.bam -I MG120.reduced.bam -I MG121.reduced.bam -I MG123.reduced.bam -I MG124.reduced.bam -I MG125.reduced.bam -I MG126.reduced.bam -I MG127.reduced.bam -I MG128.reduced.bam -I MG129.reduced.bam  -I MG130.reduced.bam -I MG131.reduced.bam -I MG132.reduced.bam -I MG133.reduced.bam -I MG134.reduced.bam -I MG135.reduced.bam -I MG136.reduced.bam -I MG138.reduced.bam -I MG139.reduced.bam -I MG140.reduced.bam -I MG141.reduced.bam -I MG142.reduced.bam -I MG143.reduced.bam -I MG144.reduced.bam -I MG145.reduced.bam -I MG146.reduced.bam -I MG149.reduced.bam -I MG150.reduced.bam -I MG151.reduced.bam -I MG152.reduced.bam -I MG154.reduced.bam -I MG155.reduced.bam -I MG156.reduced.bam --output_mode EMIT_ALL_SITES -o FinalGeno_137Lj_EAS.vcf'
}


### ----------Genetic diversity--------------###
{
# Cmd to extract few accessions :  
/com/extra/vcftools/0.1.9/bin/vcftools --keep subpop2.txt --vcf 137Lj_filt1_g5.nr.noChr0r.vcf --recode --recode-INFO-all --out 137Lj_filt1_g5.nr.noChr0r.subpop2

# Cmd for pi calculation for each chromosome in each subpopulation:
/com/extra/vcftools/0.1.14/bin/vcftools --vcf 137Lj_filt1_g5.nr.noChr0r.subpop1.recode.chr1.vcf --window-pi 62285374 --out 2_p1.chr1.out

# Normalize the pi values for the callable positions for each chromosome: Pi value * chromosome length / number of callable fraction positions
}


### ----------------PCA----------------------###
{
#Outlier accessions excluded: MG020, MG151
#Working directory:  /home/niraj/LotusGenome/Niraj/2_LotusAccessions/runGATKv2.7-2/PCA

## Generate input for the smartpca software
perl generateInputFilesForPCA.pl <vcf file> # https://github.com/ShahNiraj/JapanHistory/blob/master/generateInputFilesForPCA.pl

## create a file named '3_par' that contains the following information.
#genotypename: 3_Lj.eigenstrat.geno.txt
#snpname: 3_Lj.eigenstrat.snp.txt
#indivname: 3_Lj.eigenstrat.ind.txt
#outputformat: EIGENSTRAT
#genooutfilename: 3_Lj.eigenstrat.geno.out.txt
#snpoutfilename: 3_Lj.eigenstrat.snp.out.txt
#indoutfilename: 3_Lj.eigenstrat.ind.out.txt
#evecoutname: 3_Lj.evec.out
#evaloutname: 3_Lj.eval.out
#numoutlieriter: 0

##cmd used to run smartpca program: 
EIG6.0beta/bin/smartpca -p 3_par > 3_logfile
}


### --------------LD decay-------------------###
{
# calculate Ld using vcftools:
/com/extra/vcftools/0.1.9/bin/vcftools --vcf ../NucleotideDiversity/137Lj_filt1_g5.nr.noChr0r.vcf --geno-r2 --ld-window-bp 50000 --max-alleles 2 --min-alleles 2 --out 137Lj_filt1_g5.nr.noChr0

# extract 200000 random positions for plotting using perl script:
perl  randomPositionsForLd.pl 137Lj_filt1_g5.nr.noChr0.geno.ld > 137Lj_filt1_g5.nr.noChr0.ld50000.few.txt # https://github.com/ShahNiraj/JapanHistory/blob/master/randomPositionsForLd.pl

# plot in R
setwd("/Users/nshah/Desktop/Niraj/PopGenetics/LinkageDisequilibrium")
chr1.ld <- read.table(file="r2.few.txt",header=T)
plot(chr1.ld$MarkerDist,chr1.ld$R^2,type="p",cex=0.2,col="grey")
loe <- loess.smooth(chr1.ld$MarkerDist,chr1.ld$R^2,family="gaussian",degree=1,span=0.4)
lines(loe$x,loe$y,col="red",lwd=2)
}


### ----------Fst analysis-------------------###
{

# pack and index the raw vcf file
{
cd $vcf_dir
source /com/extra/tabix/0.2.6/load.sh
qx --no-scratch 'bgzip -c FinalGeno_137Lj_raw.vcf > Suppl_file_2_20180924.vcf.gz'
qx --no-scratch 'tabix -p vcf Suppl_file_2_20180924.vcf.gz'
}


# put a header on the filtered VCF file
{
cd $vcf_dir
head -100 FinalGeno_137Lj_raw.vcf | perl -lane 'print $_ if (m/^#/);' | perl -pe 's/MG20/MG020/g;' > vcf.header.txt
cat vcf.header.txt FinalGeno_137Lj_raw.snps.ref.filt1.g5mac0.5.nr.recode.vcf > 20180605_filt1.g5mac0.5.nr.recode.vcf
cp 20180605_filt1.g5mac0.5.nr.recode.vcf $output_dir
}

# pack and index the filtered vcf file
{
cd $output_dir
source /com/extra/tabix/0.2.6/load.sh
qx --no-scratch 'bgzip -c 20180605_filt1.g5mac0.5.nr.recode.vcf > Suppl_file_4_20180924.vcf.gz'
qx --no-scratch 'tabix -p vcf Suppl_file_4_20180924.vcf.gz'
}


## do the Fst scan
{
# the population lists were defined as 
# pop3: > 99% pop3 
# not_pop3: < 0.1% 
# pop2: > 90% pop2
# pop1: > 90% pop1

source /com/extra/vcftools/0.1.9/load.sh
cd $output_dir
qx --no-scratch 'vcftools --vcf 20180605_filt1.g5mac0.5.nr.recode.vcf --weir-fst-pop 20180622_not_pop3.txt --weir-fst-pop 20180622_pop3.txt --out 20180622_notPop3vsPop3_fst_scan.txt'

cd $output_dir
qx --no-scratch 'vcftools --vcf 20180605_filt1.g5mac0.5.nr.recode.vcf --weir-fst-pop 20180622_pop1.txt --weir-fst-pop 20180622_pop3.txt --out 20180622_Pop1vsPop3_fst_scan.txt'

cd $output_dir
qx --no-scratch 'vcftools --vcf 20180605_filt1.g5mac0.5.nr.recode.vcf --weir-fst-pop 20180622_pop2.txt --weir-fst-pop 20180622_pop3.txt --out 20180622_Pop2vsPop3_fst_scan.txt'
cd $output_dir

qx --no-scratch 'vcftools --vcf 20180605_filt1.g5mac0.5.nr.recode.vcf --weir-fst-pop 20180622_pop1.txt --weir-fst-pop 20180622_pop2.txt --out 20180622_Pop1vsPop2_fst_scan.txt'
}


# make pop1, pop2, and pop3 VCF files for viewing in IGV
{
head -n -1 vcf.header.txt > vcf.header.slices.txt
cd $output_dir
qx --no-scratch --mem 40000 bash 20180627_vcf_slicing_v2.sh
#compress and index the files
#mkdir 20180627_vcf_subsets
mv *.txt.vcf 20180627_vcf_subsets
cd 20180627_vcf_subsets
qx --no-scratch bash 20180627_tabix.sh
}


# transpose.awk script to transpose a matrix
{
{ 
    for (i=1; i<=NF; i++)  {
        a[NR,i] = $i
    }
}
NF>p { p = NF }
END {    
    for(j=1; j<=p; j++) {
        str=a[1,j]
        for(i=2; i<=NR; i++){
            str=str" "a[i,j];
        }
        print str
    }
}
}


# 20180627_vcf_slicing_v2.sh, script for making subsets of VCF files for IGV display
{
###for x in something (pop2, pop3, live, die) do the below:
y='20180605_filt1.g5mac0.5.nr.recode.vcf
perl -lane 'print $_ unless (m/##/);' $y |cut -f 1-9 > $y.first.cols.txt
perl -lane 'print $_ unless (m/##/);' $y | awk -f transpose.awk >$y.transposed.txt
array=( "20180622_pop1.txt" "20180622_pop2.txt" "20180622_pop3.txt"  "20180622_not_pop3.txt" "20171212_live.txt" "20171212_die.txt"  )
for x in "${array[@]}"
do
# transpose, get the lines matching the relevant accessions, transpose back.
grep -F -f $x $y.transposed.txt | awk -f transpose.awk | perl -pe 's/ /\t/g;' > $y.accs.txt
paste $y.first.cols.txt $y.accs.txt > $y.nohead.txt
cat vcf.header.slices.txt $y.nohead.txt > $y.$x.vcf
done
}

# 20180627_tabix.sh, prepare VCF files for IGV viewing
{
source /com/extra/tabix/0.2.6/load.sh

for y in *.txt.vcf
do
bgzip -c $y > $y.gz
done

for y in *.gz
do
tabix -p vcf $y
done
}


## Carry out a gene-by-gene analysis for Fst signals. 
{
# select a subset of the GFF file
dir='/Users/au27857/Documents/Stig_aarhus/IGV_files/ljr30'
cd $dir
perl -lane 'if ($F[2] eq "gene" && $F[1] eq "protein_coding" && abs($F[4]-$F[3])<20000 && abs($F[4]-$F[3])>200) {print $_;}' 20130802_Lj30.sorted.igv.gff3  > 20130802_Lj30.sorted.igv.protein_coding_genes.gff3


# annotate the SNPs from the Fst analysis and the gwas run using bedtools
cd $output_dir
source /com/extra/bedtools/2.16.2/load.sh
for y in *.weir.fst.bedgraph;
do nice -n 19 bedtools intersect -loj -a 20130802_Lj30.sorted.igv.protein_coding_genes.gff3 -b $y | perl -lane 'if ($F[10] != -1) {@temp=split /\;/, $F[8]; ($junk,$id) = split /\=/, $temp[0]; print "$_\t$id";}' > $y.genes.txt;
done; 


}


}


###---------fastSTRUCTURE analysis-------------###
{

### convert VCF file to str
# ~/LotusGenome/faststorage/tomomi/Data_added_201806_fastSTRUCTURE/2row_137.sh
{
#!/bin/sh
mkdir 2row_NR_137/
for i in `seq 1 1 137`
do
echo Now,column${i}!
    
    cut -f $i ForSTRUCTURE_137.txt | cut -c 1-3 | sed 's/\//\t/g' | cut -f 1 | tr '\n' '\t' > 2row_NR_137/$i.1.txt
    cut -f $i ForSTRUCTURE_137.txt | cut -c 1-3 | sed 's/\//\t/g' | cut -f 2 | tr '\n' '\t' > 2row_NR_137/$i.2.txt

    cat 2row_NR_137/$i.1.txt 2row_NR_137/n.txt 2row_NR_137/$i.2.txt 2row_NR_137/n.txt > 2row_NR_137/$i.txt
    rm 2row_NR_137/$i.1.txt 2row_NR_137/$i.2.txt

    cat 2row_NR_137/p.txt 2row_NR_137/$i.txt > 2row_NR_137/p2.txt
    mv 2row_NR_137/p2.txt 2row_NR_137/p.txt

    rm 2row_NR_137/$i.txt

#fi

done

#cat  lSNP_name.txt C_*.txt > sC.txt

#WC=`cat 2row_NR_140/C_*.txt | wc -l`

#echo 行数は$WC

mv 2row_NR_137/p.txt 2row_NR_137/2row_ForSTRUCTURE_137.txt
}


## run the fastSTRUCTURE analysis
{
#!/bin/sh

for i in `seq 1 1 8`

do

#i=1

python structure.py\
 -K $i\
 --input=2row_ForSTRUCTURE_131-7_excludeMG002MG064_headerF_2\
 --format=str\
 --output=131+7_excludeMG002MG064.K$i\
 --full

python distruct_2.py\
 -K $i\
 --input=131+7_excludeMG002MG064.K$i\
 --output=131+7_excludeMG002MG064.K$i.png\
 --title=K_$i\
 --popfile=list.131+7_excludeMG002MG064.txt

done
}


## prepare results file for analysis
{
dir='[wokring_dir]'
cd $dir
paste accession_list_136.txt fastSTRUCTURE_result_136/136_m_K3.3.meanQ > 20180612_fastSTRUCTURE_result_136.txt
less 20180612_fastSTRUCTURE_result_136.txt
}

}


###------------PSMC analysis-------------------###
{
	# subsample BAM files 
	# usage: sub_sampling.sh [subsampling rate file]
	{
	#!/bin/sh

	# [subsampling rate file] format: [Accession ID]\t[fold coverage]. Example: MG013	8.09	
	
	
	for i in `cut -f 1 $1`
	do

	p=`grep $i $1 | cut -f 2 `
	
	samtools view -s $p -b /home/tomomi/LotusGenome/100_data/124Accessions/BAMs/$i.bam\
 	> sub_sampling_bams/$i.sub_sampling.bam


	done
	}
	
	# prepare PSMC pseudo-diploid input files from BAM files 
	# # usage: make_input_201808.sh [line number for accession 1] [first line number for accession 2] [last line number for accession 2]
	# [line number] means the row of the accessions in "/home/sua/LotusGenome/faststorage/tomomi/psmc/used_accession_list/all74_path_subsampling_checked.txt", which contains full paths to the bam files
	{
	#!/bin/sh
	
	a=$1

    for b in `seq $2 1 $3`
    do

	s1=`head -n $a used_accession_list/all74_path_subsampling_checked.txt | tail -n 1 `
	s2=`head -n $b used_accession_list/all74_path_subsampling_checked.txt | tail -n 1 `

	n1=`echo $s1 | sed 's/Gifu/"\tGifu/g' | sed 's/MG/"\t/g' | cut -f 2 | awk '{print "MG"$1}'`
	n2=`echo $s2 | sed 's/Gifu/"\tGifu/g' | sed 's/MG/"\t/g' | cut -f 2 | awk '{print "MG"$1}'`
	s3=`echo $n1.$n2 `

	echo "$s3 merge and mpileup!"

	rm combinedBAMs_201808/${s3}.bam

	samtools merge combinedBAMs_201808/${s3}.bam ${s1}.bam ${s2}.bam

	samtools mpileup -C 50 -uf /home/tomomi/LotusGenome/ljr30/lj_r30.fa combinedBAMs_201808/${s3}.bam | bcftools call -c -o combinedBAMs_201808/${s3}.vcf

	echo "$s3 mpileup fasta start!"

	bcftools view combinedBAMs_201808/${s3}.vcf |\
	vcfutils.pl vcf2fq -d 5 -D 100 - |\
	gzip > ./pseudo_diploidfiles_201808/${s3}.fq.gz

	rm combinedBAMs_201808/${s3}*

    done
	}

	# Run the PSMC analysis
	# usage: run.psmc_201808.sh [last line number for accession] [number of accessions]	
	# [last line number] means the row of last accession's pseudo-diploid file in the directory, "/home/tomomi/LotusGenome/faststorage/tomomi/psmc/pseudo_diploidfiles_201808/"
	# [number of accessions] means the number of accession's pseudo-diploid files and the analyses were done independently.
	{
			
		#!/bin/sh

		

		cd ../psmc-master/

		#run the analyses for every input independently
		for i in `ls ../psmc/pseudo_diploidfiles_st3-1_true/`
		do

		#path for input file
		file=`echo $i`

		#name for result file
		name=`echo $i | sed 's/.fq.gz//g' `

		echo "${name} fq2psmcfa start!"

		#Transforms the consensus sequence into a fasta-like format
		utils/fq2psmcfa ${file} > ${name}.psmcfa

		echo "${name} psmc start!"

		#Run the PSMC, same values as the analysis for human used for the pattern of parameters (-p)
		./psmc -N25 -t15 -r5 -p "4+25*2+4+6" -o ${name}.psmc ${name}.psmcfa

		echo "${name} psmc2history.pl start!"

		#Simulates the history inferred by PSMC
		utils/psmc2history.pl ${name}.psmc | utils/history2ms.pl > ms-cmd_${name}.sh

		echo "${name} psmc_plot.pl start!"

		#Visualize the result with the options for  mutation rate per nucleotide (-u), and for number of years per generation (-g) 
		utils/psmc_plot.pl -u 7e-09 -g 1 -R ${name} ${name}.psmc
		
	}

	# get the divergence time estimates	
	{
	## all comparisons
	# source R
	source /com/extra/R/3.4.0/load.sh
	
	dir_all=''
	cd $dir_all
	mkdir out
	qx --no-scratch 'bash 20180831_divergence_times.sh'
	# "20180831_divergence_times.sh"
	{
	source /com/extra/R/3.4.0/load.sh
	for y in *.0.txt;
	do nice -n 19 cat $y | Rscript -e 'd <- read.delim("stdin", header=FALSE)' -e 'pos <- Position(function(x) x < 5,d$V2)' -e 'year <- d$V1[pos]' -e 'cat(year,"\n")' > out/$y.year.txt;
	done;
	cd out
	cat *.year.txt > 20180831_all_psmc_comb.txt
	ls *.year.txt > filenames.txt
	paste filenames.txt 20180831_all_psmc_comb.txt | perl -pe 's/\.0\.txt\.year\.txt//; s/_genomic_20130609//; s/-//g; s/MG20/MG020/g; s/\./\t/; s/\.sub_sampling//g; s/sub_sampling\.//g; s/\.rg//g; s/rg\.//g; s/a\.//g; s/\.fq\.gz//g;' > 20180831_all_psmc_comb_ids.txt
	}
	
	#less 20180831_all_psmc_comb_ids.txt
	
	# prepare file for R analysis
	perl -lane 'print "$F[1]\t$F[0]\t$F[2]";' 20180831_all_psmc_comb_ids.txt > 20180831_all_psmc_comb_ids_mirror.txt
	cat 20180831_all_psmc_comb_ids.txt 20180831_all_psmc_comb_ids_mirror.txt > 20180831_all_psmc_comb_ids_double.txt
	wc -l 20180831_all_psmc_comb_ids_double.txt
	
	}

}


###-----------------GWA------------------------###
{


### prepare the new genotype file for EMMAX analysis

## impute genotypes
qx --no-scratch -n=1 -c=1 -m=20g 'java -Xmx20g -jar /com/extra/beagle/5.0/jar-bin/beagle.16May18.771.jar gt=FinalGeno_137Lj_raw.snps.ref.filt1.g5mac0.5.nr.recode.vcf out=impute’


# perl script for converting VCF to EMMAX input format
# gwasformatchangedto012.pl, get from https://github.com/ShahNiraj/JapanHistory


{
# adjust header
dir='n'
cd $dir
perl -pe 's/MG20/mg020/; s/MG/mg/g; s/,/\t/g;' impute.hetToRef.gwas.vcf > 20180611_impute.hetToRef.gwas.tab

## tranpose, sort, retranspose
# first tranpose
qx --no-scratch --mem 10000 'awk -f transpose.awk 20180611_impute.hetToRef.gwas.tab > 20180611_impute.hetToRef.gwas.csv.tp.txt'
less -S 20180611_impute.hetToRef.gwas.csv.tp.txt
grep "Gifu" 20180611_impute.hetToRef.gwas.csv.tp.txt | less -S

# write header
head -2 20180611_impute.hetToRef.gwas.csv.tp.txt > 20180611_gwas_tp_header.txt
less -S 20180611_gwas_tp_header.txt

# sort
tail -n +3 20180611_impute.hetToRef.gwas.csv.tp.txt | sort -k 1,1 > 20180611_impute.hetToRef.gwas.csv.tp.sorted.txt
less -S 20180611_impute.hetToRef.gwas.csv.tp.sorted.txt

# add header back
cat 20180611_gwas_tp_header.txt 20180611_impute.hetToRef.gwas.csv.tp.sorted.txt > 20180611_impute.hetToRef.gwas.csv.tp.sorted.head.txt
less -S 20180611_impute.hetToRef.gwas.csv.tp.sorted.head.txt

# transpose back
qx --no-scratch --mem 10000 'awk -f transpose.awk 20180611_impute.hetToRef.gwas.csv.tp.sorted.head.txt > 20180611_impute.hetToRef.gwas.csv.tp.sorted.head.tp.txt'

# convert back to csv
perl -pe 's/ /,/g;' 20180611_impute.hetToRef.gwas.csv.tp.sorted.head.tp.txt > 20180611_impute.hetToRef.gwas.sorted.csv
less -S 20180611_impute.hetToRef.gwas.sorted.csv
# count number of commas to check
awk -F"," '{print NF}' 20180611_impute.hetToRef.gwas.sorted.csv | sort | uniq -c
}


## Run EMMAX gwas
script_dir=""
data_dir=""
refdata_dir=""
marker_file="Supplemental_file_9"
data_file="Supplemental_file_1"
prefix=""
mac=8
date=''

cd $data_dir
nohup nice -n 19 python "$script_dir"src/gwa.py -o "$prefix" -a emmax -m "$mac" -r "$data_dir""$data_file" -f "$refdata_dir""$marker_file" > $data_file.log &

}
